# Diversification of metazoan Kexin-like proprotein convertases: insights from the leech *Helobdella*

**DOI:** 10.1101/802215

**Authors:** Wei-Yu Tao, Ya-Chih Cheng, Mi Hye Song, David A. Weisblat, Dian-Han Kuo

## Abstract

Intercellular communication is quintessential for multicellularity and often mediated by secreted peptide ligands. In Metazoa, proprotein convertases are a major class of endoproteases partaking in the proteolytic processing of these ligands, which is in turn required for their signaling activities. In vertebrates, the best-studied convertase substrates are neuropeptides, peptide hormones, and members of the TGFβ/BMP-family. Each ligand is processed by a particular subset of convertases. Therefore, the diversification of convertases may have contributed to the growing complexity of cellular communication in metazoan evolution. However, proprotein convertases have not been systematically explored in Metazoa. Here, we sampled the representative metazoan genomes and established that six Kexin-like proprotein convertases were present in the last common ancestor of protostomes and deuterostomes. Among these, we identified a novel PCSKX orthologous group (OG) that was lost in vertebrates. Spiralian protosomes have, in general, maintained all six OGs. Therefore, we characterized the functional divergence of the Kexin-like OGs in the leech *Helobdella*, an experimentally tractable spiralian. Gene expression patterns suggested that PCSK1 and PCSK2 are specialized for the processing of neuropeptides and peptide hormones in bilaterians and that the newly identified PCSKX is probably functionally similar to furin and PCSK7. Finally, we showed that, distinct from the BMP morphogen in vertebrate embryos, the convertase-mediated proteolytic cleavage is not required for the short-range BMP signaling in the dorsoventral patterning of leech ectoderm. Together, our data revealed the complexity of the Kexin-like proprotein convertase gene family and their roles in generating diverse patterns of cellular communication in Metazoa.

## Introduction

Endoproteolytic processing of secreted peptides is one of the common mechanisms for the post-translational modifications that are used to activate and inactivate biomolecules in the eukaryotic cells. In endoproteolytic processing, proprotein convertases cleave the inhibitory prodomain from the precursor peptides to produce bioactive signaling ligands, receptors, and enzymes in the secretory pathway. The metazoan proprotein convertases are structurally similar to the prokaryotic Subtilisin and the yeast Kexin and are therefore named <u>P</u>roprotein <u>C</u>onvertase <u>S</u>ubtilisin/<u>Kexin</u> (PCSK) (Seidah 2011). Metazoan PCSKs are involved in a great variety of physiological and developmental processes. For example, PCSK1 (or PC1/3) and PCSK2 (or PC2) cleave inactive precursors in secretory vesicles to produce the active forms of peptide hormones and neuropeptides (Furuta, et al. 1997; Siekhaus and Fuller 1999; Zhu, et al. 2002; Husson, et al. 2006; Rhea, et al. 2010; Zhang, et al. 2010). In contrast, furin (or PCSK3), PCSK5 (or PC5/6), PCSK6 (or PACE4) and PCSK7 (or PC7) are best known for their role in the processing of TGFβ/BMP ligands, a family of signaling molecules that have multiple important developmental roles (Dubois, et al. 1995; Cui, et al. 1998; Essalmani, et al. 2008; Künnapuu, et al. 2009; Nelsen and Christian 2009), as well as a number of other secreted or transmembrane proteins (Seidah and Chrétien 1997, 1999).

Although proprotein convertases such as furin and PCSK5-7 are often viewed as “housekeeping proteins” due to their ubiquitous expression in adult tissue, they may also make significant contributions to the regulation of morphogen signals during embryonic development, since a subtle difference in the proprotein convertase-mediated cleavage of the ligand peptides can introduce a profound effect on the range and strength of TGFβ/BMP signaling and influence the shape of a morphogen gradient (Constam 2014). Intriguingly, the order in which the multiple convertase sites on the vertebrate BMP4 proprotein are cleaved can determine its signaling range, by specifying different modes of interactions between the inhibitory prodomain and the ligand peptide (Cui, et al. 2001; Degnin, et al. 2004; Tilak, et al. 2014). Moreover, proprotein convertases that are broadly expressed in adult tissue may exhibit developmentally regulated expression during embryogenesis, being specifically upregulated in cells that produce TGFβ/BMP ligands (Constam, et al. 1996; Nelsen, et al. 2005). Functional data showed that there seems to be a tissue-specific requirement for individual convertase sites on a BMP ligand (Goldman, et al. 2006; Sopory, et al. 2010; Akiyama, et al. 2012). Given that each individual convertase has distinct, preferred cleavage sites (Nelsen and Christian 2009), tissue-specific expression of convertase activity may be a critical factor in determining the differing strength and range of TGFβ/BMP signaling between tissue types.

While it has been shown that the requirement for and consequence of convertase-mediated cleavage differ between BMP ligands and between species (Fritsch, et al. 2012), the diversity in the roles of convertase-mediated cleavage in the regulation of cellular signaling has been largely unexplored. In the leech *Helobdella*, a novel form of short-range BMP signal patterns the dorsoventral axis of ectoderm (Kuo and Weisblat 2011b). Thus, to investigate the hypothesis that differential expression of selected Kexin-like proprotein convertases contributes to such a non-canonical deployment of this BMP signal, we set out to characterize the genes encoding proprotein convertases in the leech genome.

Proprotein convertases have only been studied in a small number of mostly vertebrate model species, and the pre-vertebrate evolutionary history of this gene family is largely unknown. To provide a phylogenetic framework for our studies on the leech proprotein convertases and to gain a bird’s-eye view on the evolution of this gene family in the animal kingdom, we first carried out a survey of convertase-like genes across metazoan species. Genes that are currently described as PCSKs were in fact divided into three distantly-related groups based on their distinct structural profiles. We focused on the classical Kexin-like proprotein convertases, which are responsible for processing BMP ligands. To assign individual leech convertases to specific orthologous groups (OGs), we performed a phylogenetic analysis of proprotein convertases from representative metazoan genomes. Six Kexin-like convertases were inferred to be present in the genome of the last common ancestor of protostomes and deuterostomes. Among these, we identified a novel OG named PCSKX, which was lost in the vertebrates. On the other hand, PCSK7 has a premetazoan origin, and the five other Kexin-like convertases arose from gene duplications that occurred before the protostome-deuterostome split. Additional lineage-specific gene duplications and losses further diversified the proprotein convertase gene complement among extant bilaterians.

Spiralians, including the leech *Helobdella*, have retained all six ancestral OGs of Kexin-like convertases. We analyzed the expression pattern of individual proprotein convertase genes in the adult leech. Our results suggest that, similar to other animal species, PCSK1 and PCSK2 were likely involved in the production of neurohormones and synaptic neuropeptides in the leech. Finally, we showed that none of the leech Kexin-like convertases is specifically upregulated in the BMP-producing cells and that convertase-mediated cleavage is not required for the normal signaling activity of BMP in the dorsoventral patterning of ectoderm. Therefore, the expression and functional requirement of proprotein convertases in this patterning system are very different from that in the standard BMP morphogen system as observed in vertebrate embryos. Together, our data revealed a dynamic and complex picture for the evolution of proprotein convertases in Metazoa. In line with the diversity in number and composition of proprotein convertase genes, their roles in the regulation of the peptide ligands have also undergone lineage-specific modifications. Diversification in gene repertoire and gene function of proprotein convertases is expected to play a role in the evolution of complex cellular communication systems in Metazoa.

## Results and Discussion

### Proprotein convertase-like genes in metazoan genomes

In mammals, the PCSK designation was applied to nine different genes, whose gene products are involved in the proteolytic processing of secreted proteins (Seidah 2011). Among these, PCSK1-7 are characterized by a P domain (Pfam PF01483) on the C-terminal side of the catalytic peptidase S8 domain (Pfam PF00082) and are structurally similar to the yeast Kexin. Homologs of Kexin-like convertases, such as PCSK2/PC2 and PCSK3/furin, have been identified and characterized in a number of invertebrate species (Fan and Nagle 1996; Agata, et al. 1998; Oliva, et al. 2000; Thacker and Rose 2000; Rayburn, et al. 2003; Bertrand, et al. 2006; Cummins, et al. 2009). Published records for homologs of PCSK8 and PCSK9 outside of vertebrates are sparse, but their phylogenetic distribution suggests an ancient, pan-eukaryotic origin for each of these genes. PCSK8 is also known as Site-1 Protease (S1P), or Subtilisin/Kexin-Isozyme 1 (SKI-1); its homologs were reported in *Drosophila* (Seegmiller, et al. 2002) and in *Arabidopsis* (Liu, et al. 2007) but not in yeast. PCSK9, or Neural Apoptosis-Regulated Convertase 1 (NARC-1), is structurally similar to the fungi proteinase Ks (Naureckiene, et al. 2003; Seidah, et al. 2003); its homologs were reported in a few metazoan species, including amphioxus, sea cucumber, and gastropods (Cameron, et al. 2008; Ren, et al. 2016). Here, we will use “Kexin-like,” “S1P-like,” and “PK-like” to refer to the convertases that are structurally similar to vertebrate PCSK1-7, PCSK8, and PCSK9 respectively, for reasons entailed later in this section.

To further explore the evolution of the proprotein convertase genes in Metazoa, we searched for the Kexin-like, S1P-like, and PK-like convertase genes in selected genomes represented in the ENSEMBL databases. A Kexin-like convertase gene was defined by having an amino acid sequence that matches both Pfam PF00082 and Pfam PF01483 profiles. S1P-like convertase genes were defined by matching both Pfam PF00082 and InterPro IPR034185 (Site-1 peptidase catalytic domain). PK-like convertase genes were defined by matching all of the following three profiles, namely Pfam PF00082, Pfam PF05922 (Peptidase inhibitor I9) and InterPro IPR034193 (Proteinase K-like catalytic domain). Gene models that match any one, but not all, of the defining profiles were manually examined and analyzed by reciprocal BLAST search to identify incomplete models and to remove the false positives. The complete set of convertase genes identified in this study is listed in Table S1.

Among the three non-metazoan opisthokont genomes searched, one Kexin-like gene (*KEX2*) and three S1P-like genes (*RRT12*, *PRB1*, and *YSP3*) were found in the yeast *Saccharomyces cerevisiae*; one Kexin-like, one S1P-like and three PK-like genes were found in the filopodia-bearing, amoeboid protist *Capsaspora owczarzaki*; and two Kexin-like genes and one S1P-like were found in the choanoflagellate *Monosiga brevicollis*. We also searched the genome of *Acanthamoeba castellanii*, and found three Kexin-like, one S1P-like, and nine PK-like genes. *A. castellanii* belongs to Amoebozoa, a distant outgroup for Opisthokonta (Cavalier-Smith, et al. 2014).

Within Metazoa, we searched 23 genomes, including five vertebrates (the human *Homo sapiens*, the chicken *Gallus gallus*, the western clawed frog *Xenopus tropicalis*, the zebrafish *Danio rerio*, and the lamprey *Petromyzon marinus*), three invertebrate deuterostomes (the ascidian *Ciona intestinalis*, the amphioxus *Branchiostoma floridae*, and the sea urchin *Strongylocentrotus purpuratus*), four ecdysozoans (the fruit fly *Drosophila melanogaster*, the red flour beetle *Tribolium castaneum*, the water flea *Daphnia pulex*, and the nematode *Caenorhabditis elegans*), seven spiralians (the limpet *Lottia gigantea*, the oyster *Crassostrea gigas*, the octopus *Octopus bimaculoides*, the polychaete worm *Capitella teleta*, the leech *Helobdella robusta*, the brachiopod *Lingula anatina*, and the rotifer *Adineta vaga*), and four non-bilaterians (the sea anemone *Nematostella vectensis*, the placozoan *Trichoplax adherens*, the ctenophore *Mnemiopsis leidyi*, and the sponge *Amphimedon queenslandica*). The copy number of Kexin-like genes in these metazoan genomes ranged from three (*Trichoplax*) to nineteen (*Adineta*) and was never zero in any of these species (Figure 1A). The copy number of S1P-like convertases ranged from zero (*Caenorhabditis*) to three (*Lingula*); a single copy was found in 19 out of the 23 metazoan genomes (Figure 1A). The PK-like convertase genes were absent from 15 out of the 23 genomes and were highly duplicated in others, with the most extreme case being the sea urchin genome which contained as many as unique 62 PK-like loci (Figure 1A).

**Figure 1.**
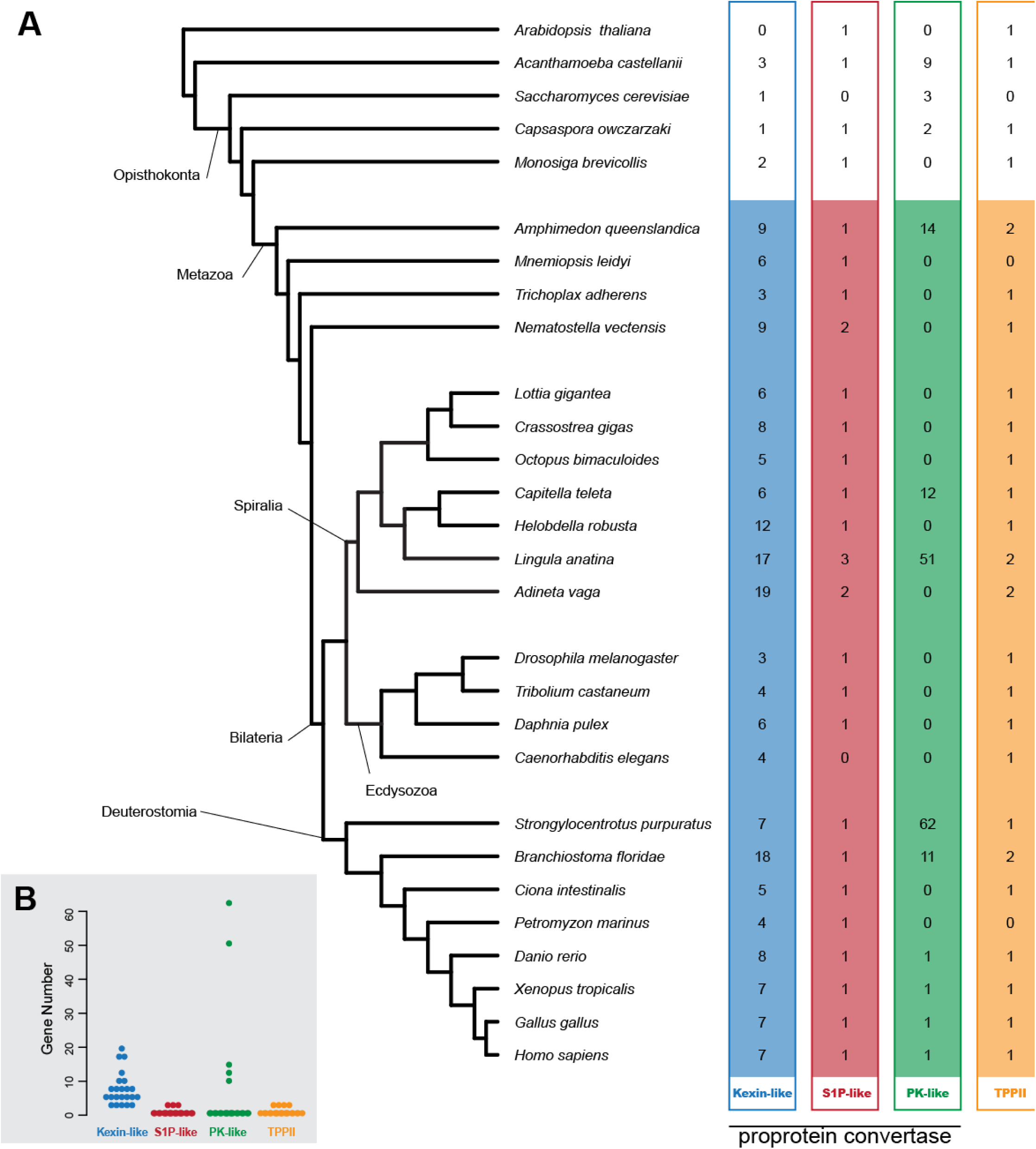
Phylogenetic distribution of metazoan S8 domain-containing peptidases. **A.** The number of Kexin-like, S1P-like, PK-like, and TPPII genes in 23 metazoan and five non-metazoan genomes analyzed in this study. **B.** Beeswarm plot showing the variability in the gene family size of Kexin-like (blue dots), S1P-like (red dots), PK-like (green dots), and TPP (orange dots) in individual metazoan genomes. Each dot represents the number of genes in the specified gene family in a metazoan genome.

A fourth, and the final, group of genes encoding S8 domain-containing peptidases, namely the tripeptidyl peptidase II (TPPII) family, was also broadly represented in the metazoan genomes. TPPII functions in cytosolic proteolysis (Rockel, et al. 2012), but not in the secretory pathway. By definition, TPPII is not a proprotein convertase. For the sake of completeness, we nevertheless went on to document TPPII-like genes in the metazoan genomes. Genes of this family can be structurally defined by matching Pfam PF00082 and Pfam PF12580 (TPPII). The copy number of TPPII-like genes in the 23 metazoan genomes examined ranged between zero and two. TPPII-like was absent from the genomes of *Mnemiopsis* and *Petromyzon*. Two copies of TPPII-like genes were found in *Amphimedon*, *Lingula*, *Adineta*, and *Branchiostoma*. In the remaining 17 genomes, each contained only one TPPII-like gene (Figure 1A). Outside of Metazoa, single TPPII-like genes were found in the genomes of *Acanthamoeba*, *Capsaspora*, and *Monosiga,* but not in the yeast *Saccharomyces*, suggesting a gene loss in the lineage leading to this species. Overall, the copy number of S1P-like and TPPII-like genes showed little variation between metazoan genomes, whereas the copy number variation was greater for the Kexin-like and the PK-like genes (Figure 1B)

In addition to the four structurally defined S8-like peptidase gene families we found in metazoan genomes, Kexin-like, S1P-like, PK-like, and TPPII-like, a more diverse repertoire of peptidase S8-like genes was uncovered outside of Metazoa (Bryant, et al. 2009; Li, et al. 2017). Based on the following observations, we postulated that the four metazoan S8-like families are probably distantly related and not evolutionary sisters. First, from their distribution across species, each of the four families has a very deep origin, probably pre-Eukaryota. Second, each family has a distinct structural profile from each other and recognizes a distinct set of target-site sequences (Seidah, et al. 1999; Seidah, et al. 2003; Seidah 2011). Third, sequences of members from different families cannot be aligned without catastrophic loss of phylogenetic information (data not shown). Given that TPPII-like does not have a proprotein convertase function and that the four families of S8-like genes in the metazoan genomes are distantly related, we speculate that the propeptidase function of the three families of S8-like peptidases arose by functional convergence. Thus, although PCSK has been used to refer to the metazoan S8 peptidase genes possessing a proprotein convertase activity, one should refrain from using PCSK for the larger gene families on a phylogenetic basis; we therefore use specific names such as Kexin-like, S1P-like, and PK-like to refer to the gene families here.

### Kexin-like proprotein convertase genes in the *Helobdella* genome

Pfam query and BLAST search of the *Helobdella robusta* database yielded 17 candidate gene models encoding Kexin-like genes. Among these, nine models were determined as being complete or nearly complete, based on the criterion that they have all the structure domains that are typical of known Kexin-like convertases, namely a prodomain, a catalytic domain, and a P domain. The remaining eight are apparently incomplete models. We next examined the details of these incomplete models to revise these models. By manually examining the genome location of these models, we found that three models (JGI Protein IDs: 74170, 137839, and 149650) are adjacent to each other and that each corresponds to a unique region of a typical Kexin-like convertase. Therefore, we propose that these three models should be merged to represent a single gene in the *Helobdella* genome. Another two models (JGI Protein IDs: 81814 and 184031) each represent a different portion of a typical Kexin-like convertase, and each is located at the end of a large scaffold. Therefore, we proposed that these two models may represent a single gene and that they were separated due to incomplete genome assembly. Third, one gene model (JGI Protein ID: 190143) contains a gap in the sequence between two predicted exons, and two models (JGI Protein IDs: 93024 and 93071) are located on small scaffolds, 1,371 and 1,142 base-pairs in length respectively. Of these two small scaffolds, gene model 93024 fits into the gap in gene model 190143; thus, we proposed that these two models also represent a single gene. Finally, the sequence of the genome scaffold containing the remaining orphaned model (JGI Protein ID: 93071) was derived from two relatively short Sanger reads with poor quality and is highly similar to a small region on Scaffold 34 that contains model 82977. Therefore, we determined that model 93071 and the genome scaffold containing it (scaffold 3727) are probably artifacts resulting from poor sequencing reads. Together, our *in silico* approach identified 12 candidate Kexin-like convertase genes in the *Helobdella* genome (Table S1). These predictions were confirmed by cloning and sequencing of cDNA for each of the 12 predicted genes. A single S1P-like (JGI protein ID: 194246) and a single TPPII-like (JGI protein ID: 107485) were identified in the *Helobdella* genome. No PK-like gene was found.

### Phylogeny of metazoan Kexin-like genes

To assign orthologies to the 12 *Helobdella* Kexin-like convertases, we next performed a phylogenetic analysis. From the gene list uncovered by Pfam profile search, gene models that contain only partial sequence, as determined by whether all the Pfam-defined domains are completely covered by the model sequence, were excluded from our analysis. As a result, all four *Petromyzon* models, and three *Amphimedon*, one *Nematostella*, one *Crassostrea*, one *Octopus*, five *Lingula*, three *Adineta*, and two *Strongylocentrotus* models were removed from the dataset. Furthermore, given the high degree of conservation among gnathostome Kexin-like gene complements, only three vertebrate species (*H. sapiens*, *G. gallus*, and *X. tropicalis*) were included in our analysis to reduce computation load. The 12 *Helobdella* Kexin-like protein sequences were revised based on the cloned cDNA sequences.

The amino acid sequence alignment of the 156 genes from the genomes of three non-metazoan opisthokonts and 21 metazoans was produced, and a Maximum-Likelihood (ML) tree was then generated by applying LG+G model to the dataset, with *Saccharomyces* Kexin as an outgroup. In this tree, four well supported OGs were recognized: PCSK7, PCSK1, PCSK2, and a novel OG consisting entirely of previously uncharacterized invertebrate proprotein convertase genes (Figure 2A). We named this novel OG “PCSKX.” One *Helobdella* ortholog was found in each of the four OGs. We therefore named these *Helobdella* genes *Hro-pcsk7*, *Hro-pcsk1*, *Hro-pcsk2,* and *Hro-pcskx*. Among the remaining eight *Helobdella* Kexin-like genes, six were in two poorly supported groups clustered with the vertebrate furin/PCSK4 genes and protostome furin1 genes, and two were in groups consisting of apparently long-branch genes.

**Figure 2.**
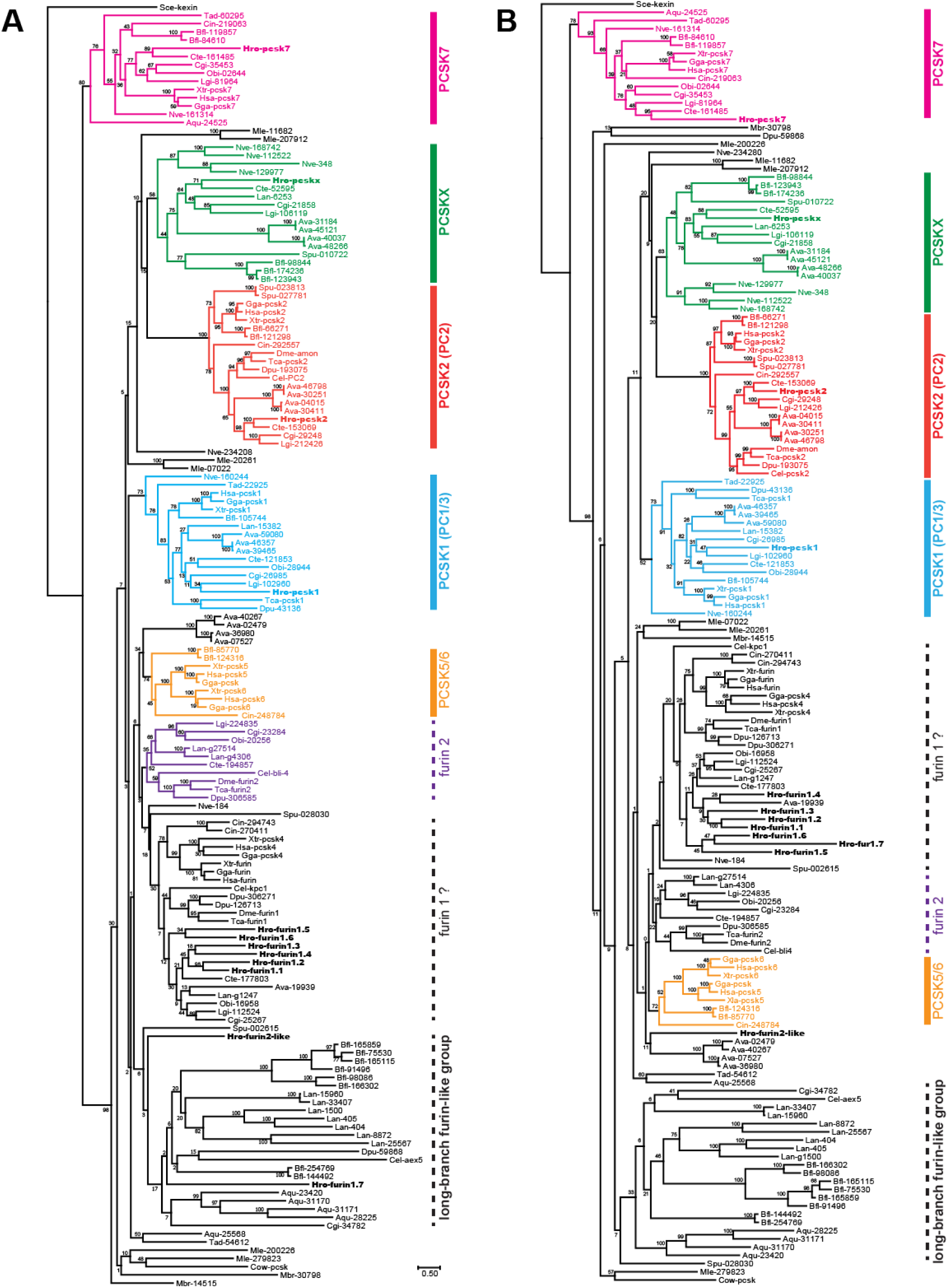
Phylogenetic analysis of the 156 Kexin-like proteins from 21 metazoan and three opisthokont genomes. **A.** The best maximum-likelihood tree produced by using the LG model. **B.** The best maximum-likelihood tree produced by using the CAT-GTR model. 500X bootstraps were performed for each tree. The three-letter prefix in the taxon name indicates the species from which the protein was identified. The amino acid sequences retrieved from the genome of *Helobdella robusta* were labeled in bold types. See Table S1 for the details of the taxon label. The thick vertical bars on the right side of the tree mark better supported OG, whereas the dashed line denotes a poorly supported OG. The branching order of genes in the better supported OGs is generally in agreement with the species tree.

We hypothesized that such a tree topology might result from a long-branch attraction (LBA) artifact. We next sought to eliminate the potential LBA by using a 77-gene/15-species dataset, in which species other than *Helobdella* containing duplicated, long-branch genes were removed. Besides *Helobdella*, each of the remaining ten bilaterian species in this dataset has at most seven Kexin-like genes in its genome. In the tree constructed under LG+G model, seven of the *Helobdella* genes were in a clade that also includes protostome furin1 genes and vertebrate furin/PCSK4 genes. We therefore named these seven *Helobdella* furin1 paralogs *Hro-furin1.1*, *Hro-furin1.2*, *Hro-furin1.3*, *Hro-furin1.4*, *Hro-furin1.5*, *Hro-furin1.6*, and *Hro-furin1.7* respectively (Figure 3A). The remaining *Helobdella* gene was in a poorly supported group that includes the protostome furin2 genes and a sea urchin gene (Figure 3A). This *Helobdella* gene is named *Hro-furin2-like*. Hence, the removal of long-branch genes had apparently improved the phylogenetic resolution to allow us to assign each of the 12 *Helobdella* Kexin-like genes to a specific OG. By analyzing the complete 156-gene/24-species dataset using the CAT-GTR model, which generally outperforms other models in resolving long branches (Stamatakis 2006; Lartillot, et al. 2007; Price, et al. 2010; Izquierdo-Carrasco, et al. 2011), we obtained a similar grouping of the *Helobdella* furin-like genes (Figure 2B). The fact that similar results were obtained using two different approaches gives us confidence that the orthology assignments for the *Helobdella* furin-like genes are likely to reflect their true phylogenetic affiliations.

**Figure 3.**
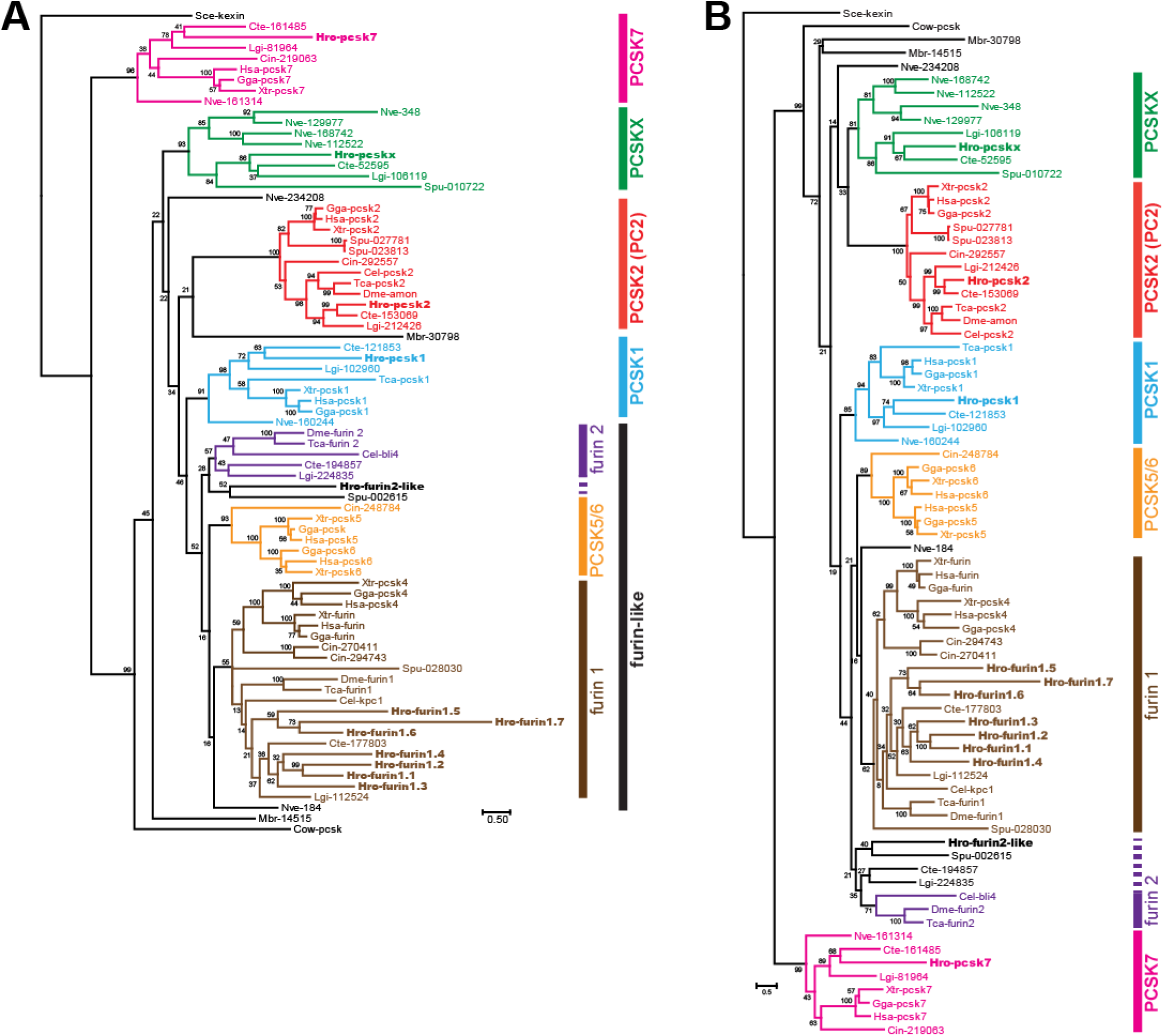
Assignment of the leech Kexin-like genes to specific orthologous groups (OGs) based on phylogenetic trees of the 77 proteins from 12 metazoan and three opisthokont genomes. **A.** The best maximum-likelihood tree produced by using the LG+G model. **B.** The best maximum-likelihood tree produced by using the CAT-GTR model. 500X bootstraps were performed for each tree. The three-letter prefix in the taxon name indicates the species from which the protein was identified (refer to Figure 1). The amino acid sequences retrieved from the genome of *Helobdella robusta* (Hro) are labeled in bold type. See Table S1 for the details of the taxon label. The thick vertical bar on the right side of the tree marks a well supported OG.

The placement of one of the choanoflagellate Kexin genes in the 77-gene/15-species LG+G tree is inconsistent with the currently accepted phylogenetic hypothesis. We next used the CAT-GTR model to analyze the 77-gene/15-species dataset. In the resulting tree (Figure 3B), the first node from the root separates PCSK7 from the other Kexin-like genes. On the other side, the *Capsaspora* gene branches off first, followed by a group consisting of the two *Monosiga* paralogs. Although these nodes are not well supported, they are consistent with the currently accepted species tree. The grouping of PCSK1, PCSK2, PCSKX, furin1, furin2 (containing non-chordate species only), and PCSK5/6 (containing chordate species only) was identical between LG+G and CAT-GTR trees, but the branching order differs. In both cases, PCSK1 and the large furin-like group (furin1 + furin2 + PCSK5/6) are sister clades. In the CAT-GTR tree, PCSK2 and PCSKX are sister clades, and the (PCSK2 + PCSKX) clade is, in turn, the sister of the (PCSK1 + furin-like) clade. In the LG+G tree, PCSK2 is the sister of the (PCSK1 + furin-like) clade, and PCSKX is the sister of the (PCSK2 + (PCSK1 + furin-like)) clade. However, none of these deep nodes is well-supported.

To further resolve the interrelationships between the identified OGs, we next removed the *Helobdella* genes, which also represent long branches themselves, and performed the ML tree search using either LG+G or CAT-GTR model. In LG+G trees as well as CAT-GTR trees, the interrelationships between PCSKX, PCSK1, PCSK2, and furin-like groups were the same regardless of whether the *Helobdella* genes were included or not (Figure 3A *versus* Figure 4A; Figure 3B *versus* Figure 4B). When polytomy was allowed for conditions in which the bootstrap support was < 50%, the topologies of LG+G tree and CAT-GTR tree were very similar to each other, with only one difference detected: the separation of the two sea urchin furin-like genes into the two furin-like subgroups in the CAT-GTR tree (*c*.*f*. Figure 4A’ and Figure 4B’). The support for the sisterhood between PCSK1 and furin-like genes was nevertheless improved in the CAT-GTR tree. Notably, this CAT-GTR tree (Figure 4B) provided a plausible parsimonious solution to the phylogeny of the furin-like genes. In the moderately supported furin-like group, a single *Nematostella* gene branched off first; next, a well-supported furin1 subgroup and a poorly supported (furin2 + PCSK5/6) subgroup separated. This branching pattern provided the best solution to reconcile the species tree. Therefore, we proposed the (furin2 + PCSK5/6) grouping as our working phylogenetic hypothesis.

**Figure 4.**
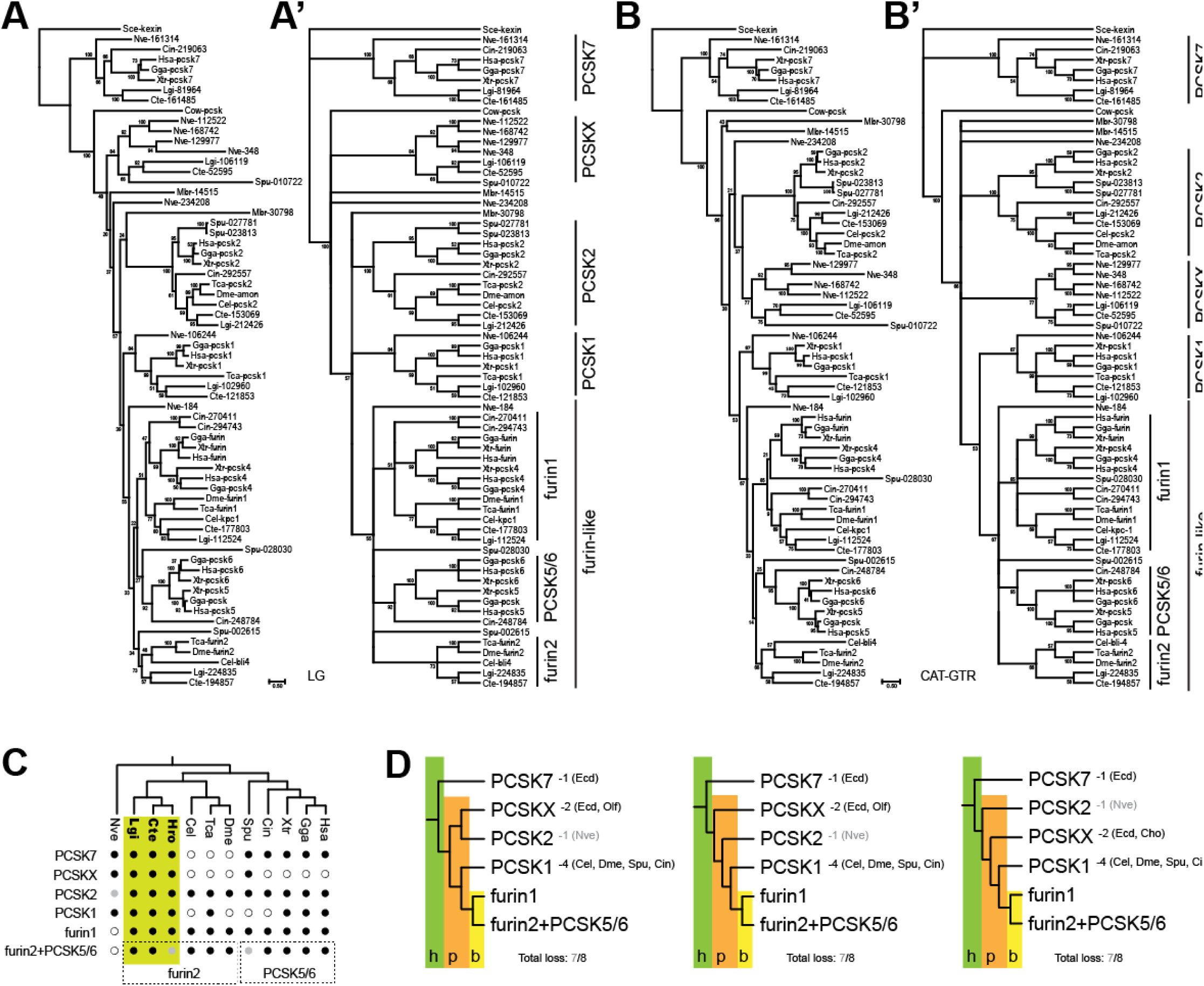
Phylogenetic analysis of the 65 Kexin-like protein from the 11 metazoan species that have less divergent Kexin-like gene complements and the three opisthokont outgroups revealed the evolutionary history of metazoan Kexin-like genes. **A.** The best maximum-likelihood tree produced by using the LG model. **B.** The best maximum-likelihood tree produced by using the CAT-GTR model. 500X bootstraps were performed for each tree. Panels A’ and B’ show cladograms of the trees in panels A and B respectively, but with nodes having less than .50 bootstrap support collapsed and shown as polytomies. **C.** The pattern of gene loss in the metazoan Kexin-like gene family. A closed black circle indicates the presence of identified ortholog(s) in a genome. An open circle denotes the absence of specific ortholog in a genome. A grey closed circle denotes the potential existence of derived homolog(s) in a genome (see text for details). Note that spiralian species (yellow) contains all six OGs. **D.** Three alternative phylogenetic scenarios which all entail equal numbers of gene losses. Ecd: gene loss in the common ancestor of ecdysozoans; Ver: gene loss in the common ancestor of Olfactores (Vertebrata + Urochordata); Nve: gene loss in *Nematosella vectensis*; Cel: gene loss in *Caenorhabditis elegens*; Dme: gene loss in *Drosophila melanogaster*; Spu: gene loss in Strongylocentrotus purpuratus; Cin: gene loss in *Ciona intestinalis*. The green shade marked with “h” indicates that the gene duplication event giving rise to PCSK7 and the ancestor of the remaining five OGs was premetazoan, probably in a holozoan ancestor. The orange shade marked with “p” indicates gene duplication events in the planulozoan ancestor, from which the modern-day PCSKX, PCSK2, and PCSK1 orthologous group arose. The yellow shade marked with “b” indicates the gene duplication event that gave rise to furin1 and furin2+PCSK5/6 OGs in the bilaterian ancestor.

We next attempted to infer the evolutionary history for each of the six OGs by reconciling our gene trees to the species tree. The species tree used here was synthesized from recent phylogenomic studies (Laumer, et al. 2015; Telford, et al. 2015; Feuda, et al. 2017; Kocot, et al. 2017). PCSK7 was identified in many sampled metazoan genomes, including the basally branching species such as *Amphimedon*, *Trichoplax*, and *Nematostella* (Figure 2, Figure 4C). Although the sea urchin PCSK7 gene was missing from our phylogenetic analysis, a reciprocal BLAST search identified one of the two incomplete *Strongylocentrotus* gene models that were dropped from the dataset as being most similar to PCSK7, suggesting that PCSK7 is likely present in this species. Given that all the *Capsaspora* and *Monosiga* Kexin-like genes branch off after the PCSK7 group, PCSK7 might have arisen by a gene duplication event in the ancestral filosozoan or holozoan, followed by gene losses in lineages such as *Capsaspora* and *Monosiga*. Within Metazoa, PCSK7 was apparently lost in the ecdysozoan lineage.

The PCSKX is a novel Kexin-like OG identified in this study, with members present in *Nematostella*, several spiralian species, sea urchin, and amphioxus, but not in ecdysozoans and vertebrates (Figure 2, Figure 4C). This suggests two independent gene losses in these respective lineages. This OG was likely to have arisen before the common ancestor of planulozoans, the group consisting of cnidarians and bilaterians. Similar to PCSKX, the PCSK1 orthology group contains genes from *Nematostella* and bilaterians, and therefore, the PCSK1 OG likely predated the planulozoans, and was lost independently in some bilaterian species, *e*.*g*. *Caenorhabditis*, *Drosophila*, *Strongylocentrotus*, and *Ciona*.

We could not robustly resolve the phylogenetic relationships among the furin-like subgroups. Nevertheless, three better-supported subgroups can be identified: furin1 (present across all bilaterians; including the vertebrate furin/PCSK4), furin2 (in protostomes only), PCSK5/6 (in vertebrate only). If we accept the (furin2 + PCSK5/6) grouping, a simple scenario for the evolution of furin-like genes emerges: first, a gene duplication took place in the ancestral planulozoan and gave rise to a PCSK1 gene and a furin-like gene. Next, in the ancestral bilaterian, the furin-like gene duplicated into furin1 and furin2. In vertebrates, furin1 further duplicated into furin (a.k.a. PCSK3) and PCSK4, while furin2 duplicated into PCSK5 and PCSK6. In a more complicated alternative scenario, a series of duplications from the planulozoan furin-like gene led to three genes, furin1, furin2, and PCSK5/6, in the ancestral bilaterian. Furin1 was preserved in all bilaterian lineages. In contrast, furin2 was lost in the chordates, and PCSK5/6 was lost in protostomes.

PCSK2 was found across bilaterians but not in any non-bilaterian metazoan. This may lead one to conclude that PCSK2 originated in Urbilateria. However, this is inconsistent with the species represented in its candidate sister groups, *i.e.* the PCSKX, PCSK1, and furin-like OGs. Since these all contain genes from non-bilaterian metazoan species, even by considering the phylogenetic uncertainty between these OGs, none of the possible tree topologies is consistent with an urbilaterian origin of the PCSK2 group. To reconcile this discrepancy, we hypothesize that PCSK2 originated before the ancestral planulozoan and was lost in the lineage leading to *Nematostella*. We have not found PCSK2 in the genomes of the hydra and the coral *Acropora*, suggesting that the loss of PCSK2 probably took place in the common ancestor of modern cnidarians. However, an orphaned *Nematostella* gene, Nve-234208, tended to lie near PCSK2 in our phylogenetic analyses and may be in fact a highly derived PCSK2. Orthologs of Nve-234208 were also identified by reciprocal BLAST in the genomes of hydra and coral, suggesting that this is not an anomaly unique to *Nematostella*. In that case, the cnidarian PCSK2 may have undergone a rapid sequence divergence instead of a gene loss.

We could not reach a single consensus conclusion on the interrelationships between the bilaterian Kexin-like OGs. Our data provide soft support for the sisterhood between PCSK1 and furin-like OGs. By constraining them as sister groups, there were three possible scenarios (Figures 4D), all of which were equal in the number of gene losses required to reconcile the species tree. In our phylogenetic analyses, however, PCSK1 was most likely the sister of the furin-like genes, and therefore, the phylogenetic uncertainty mostly lies in the branching order of PCSK2, PCSKX, and the (PCSK1 + furin-like) clade.

In summary, we propose that the last common ancestor of modern bilaterians had six Kexin-like genes, namely PCSK7, PCSKX, PCSK2, PCSK1, furin1, and (furin2+PCKS5/6). We have identified an early origin of PCSK7 but not been able to fully resolve the details in the early history of Kexin-like genes. Trees that include genes from the most basally branching metazoans, *Amphimedon* and *Mnemiopsis*, could not be reasonably reconciled, and this indicates a skewed tree topology, presumably resulting from LBA. We also find that taxon-specific duplication events produced further local diversification of selected Kexin-like genes. For example, both the leech *Helobdella* and the brachiopod *Lingula* have multiple furin-like genes. It is interesting to note the bdelloid rotifer *Adineta* has four paralogs for each of PCSKX, PCSK1, PCSK2, and furin2, consistent with the degenerate tetraploid genome that is characteristic of this asexual group (Hur, et al. 2009; Flot, et al. 2013; Nowell, et al. 2018). Furthermore, although amphioxus has been inferred as having a genome organization representative of the ancestral chordate (Putnam, et al. 2008), the amphioxus Kexin-like genes are unexpectedly highly duplicated, as multiple paralogs of PCSK7, PCSKX, PCSK2, and furin-like were found in this species (Figure 2). On the other hand, selected OGs have been lost in specific taxa. For example, ecdysozoans have lost PCSK7 and PCSKX, and the vertebrates have lost PCSKX. From our dataset, it seems obvious that the spiralian species have maintained a complete set of the six Kexin-like genes that were likely present in the last common ancestor of bilaterians (Figure 4C). Although we did not succeed in identifying the amphioxus furin1 ortholog(s), we could not exclude that they are in fact represented by the highly duplicated long-branch furin-like genes (Figure 2). In that case, the amphioxus genome would also contain all six OGs since clear-cut orthologs of PCSK1, PCSK2, (furin2+PCSK5/6), PCSK7, and PCSKX were identified in amphioxus, and we expect that the presence of all six OGs should be prevalent in the basally branching deuterostome taxa, such as echinoderms and hemichordates.

### Functional divergence of proprotein convertase genes

The fact that the six Kexin-like orthology groups were already in place in Urbilateria, and have been conserved to varying degrees since that time is indicative of a certain degree of functional divergence, which is expected to facilitate the preservation of duplicated genes (Ohno 1970). Consistent with this, the mammalian Kexin-like convertases are not functionally redundant. PCSK1 and PCSK2 are involved in the processing of neuropeptides and hormones as they are expressed in brain, neuroendocrine and endocrine tissues, but their null mutant phenotypes differ (Smeekens, et al. 1991; Furuta, et al. 1997; Zhu, et al. 2002; Refaie, et al. 2012); PCSK4 is expressed in the reproductive system and is required for fertility (Seidah, et al. 1992; Mbikay, et al. 1997; Tadros, et al. 2001; Qiu, et al. 2005); furin and PCSK5-7 are broadly expressed, but these genes also exhibit distinct mutant phenotypes (Roebroek, et al. 1998; Constam and Robertson 2000; Thomas 2002; Essalmani, et al. 2006; Wetsel, et al. 2013).

We hypothesize that the origin of functional specialization observed among the mammalian proprotein convertases can be traced back to the bilaterian ancestor and will thus show parallels to modern representatives of distantly related taxa. This hypothesis would predict (1) that the expression of PCSK1 and PCSK2 should be restricted to the nervous system and/or endocrine system across Bilateria, and (2) that furin1, furin2 and PCSK7 should be broadly expressed in most bilaterian species. The newly discovered PCSKX group has obviously not been characterized, and therefore, new expression data will help to understand its biological function. Having confirmed that the spiralians, in general, have all six Kexin-like OGs, gene expression data from a spiralian species is well suited for testing our hypothesis. Therefore, we characterized expression patterns of the 12 Kexin-like convertases in differentiated tissues by whole-mount *in situ* hybridization (WMISH) to juvenile specimens of the leech *Helobdella austinensis*, an experimentally tractable spiralian species (Weisblat and Kuo 2009, 2014).

#### Tissue-specific expression of PCSK1 and PCSK2

As predicted, both *Hau-pcsk1* and *Hau-pcsk2* exhibited tissue-restricted expression in the juvenile leech. *Hau-pcsk1* transcript was prominently expressed in a few pairs of cells in the supraoesophageal ganglion (Figure 5A). In ganglia of the ventral nerve cord, however, lower levels of Hau-pcsk1 expression were detected in a specific set of cells (Figure 5B). In contrast, *Hau-pcsk2* was broadly expressed in the central nervous system (CNS), including the ventral nerve cord and supraoesophageal ganglion (Figure 5C, 5D). Therefore, *Hau-pcsk1* has a more restricted expression than *Hau-pcsk2* in the CNS. Outside of the CNS, *Hau-pcsk2* is also expressed in clusters of large cells flanking the gut on the dorsal side (Figure 4C-4E). Based on their sizes and positions, we speculated that these cells are likely not neurons but exocrine cells of the salivary glands (Moser and Desser 1995; Kuo and Weisblat 2011a).

**Figure 5.**
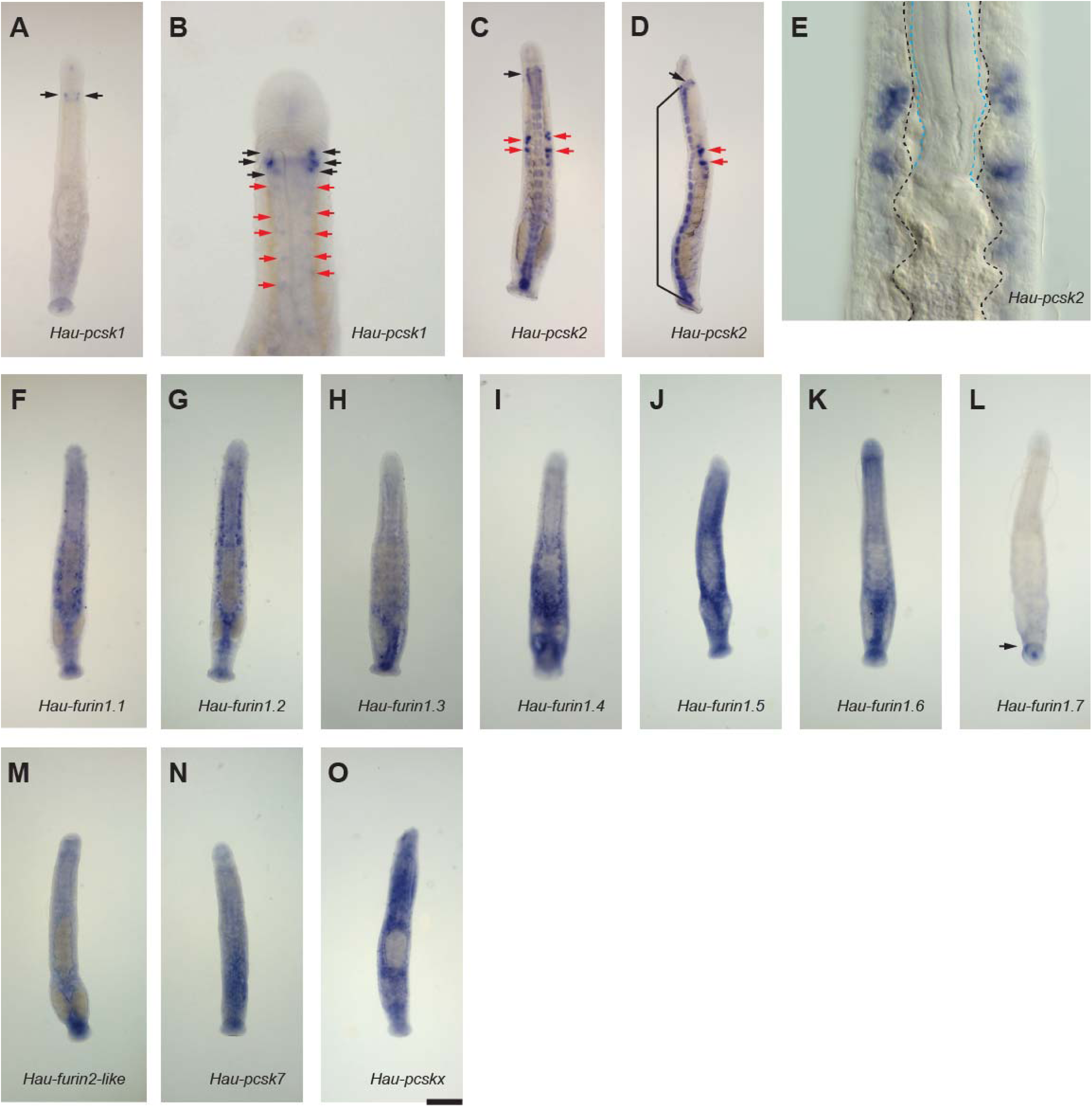
Expression patterns of the Kexin-like genes in juvenile *Helobdella austinensis* revealed by whole-mount *in situ* hybridization. *Hau-pcsk1* (A and B) is prominently expressed in paired cell clusters in the supraoesophageal ganglion (black arrows). However, lower levels of *Hau-pcsk1* transcript were also detected in cells of the ventral segmental ganglia (red arrows in B). *Hau-pcsk2* (C-E) is broadly expressed in the central nervous system, consisting of the supraoesophageal ganglion (black arrows in C and D) and the 32 ventral segmental ganglia (bracket in D). In addition to the neural tissue, *Hau-pcsk2* was also expressed in cells of the paired salivary glands (red arrows in C and D). High-power view (E) shows that the large cells expressing *Hau-pcsk2* are adjacent to the wall of the digestive tract (black dashed lines). The retracted proboscis is marked with blue dashed lines. *Hau-furin1.1* (F), *Hau-furin1.2* (G), *Hau-furin1.3* (H), *Hau-furin1.4* (I), *Hau-furin1.5* (J), *Hau-furin1.6* (K), *Hau-furin2-like* (M), *Hau-pcsk7* (N) and *Hau-pcskx* (O) were broadly expressed in juvenile leech, although sites of elevated gene expression differ somewhat among these genes. In contrast, *Hau-furin1.7* (L) was specifically expressed in the rear sucker (black arrow). Panels A-B and E-O are the dorsal view; panel C shows the ventral view; panel D shows the lateral view. Scale bar = 150 μm in panels A, C, D, and F-O; 75 μm in panel B; 30 μm in E.

Invertebrate PCSK1 orthologs have not been characterized in species other than *Aplysia* (Gorham, et al. 1996; Ouimet and Castellucci 1997) and *Helobdella* (this study). In *Aplysia*, the PCSK1 homolog is expressed in the ganglionic neuroendocrine cells that produce the reproductive hormone (Ouimet and Castellucci 1997) as well as the exocrine atrial gland that produces the reproductive pheromone (Gorham, et al. 1996). Our data showed that the leech PCSK1 is prominently expressed in the supraoesophageal ganglion, representing the brain of the leech. Given that many invertebrate peptide hormones, including those of the leech, are produced by neurosecretory cells of the brain (Grimmelikhuijzen, et al. 1980; Engelhardt, et al. 1982; Vigna, et al. 1984; Pestarino 1990; van Heumen and Roubos 1990; Gomot, et al. 1992; Meester, et al. 1992; Sonetti, et al. 1992; Kellner-Cousin, et al. 1994; Kellner, et al. 2002; Wang, et al. 2007; Miller and Newmark 2012), these specialized *Hau-pcsk1*-expressing cells may well be some of the neurosecretory cells of the leech.

On the other hand, *Hau-pcsk2* is broadly expressed in the CNS, and this is similar to the expression pattern of planarian PCSK2 (Shimoyama, et al. 2016). Not all invertebrate PCSK2 genes exhibit broad CNS expression, however. For example, the expression of *Drosophila* PCSK2 seems to be restricted to a small subset of neurons and endocrine cells (Siekhaus and Fuller 1999). Such a difference may be indicative of the varying abundance of peptidergic neurons among these species. Alternatively, it may signify the evolution of other neural functions than neuropeptide processing for PCSK2s in leech and planarian (Lophotrochozoa) compared to *Drosophila* (Ecdysozoa). It is noteworthy that, in addition to the neural tissue, PCSK2 was shown to be expressed in the atrial gland, an exocrine gland, in *Aplysia* (Nagle, et al. 1995). This seems to parallel the expression of *Hau-pcsk2* in the salivary gland. We interpreted these as independent recruitment of PCSK2 expression to the leech salivary and *Aplysia* atrial gland, however, since the two glands are not homologous. In any case, our data were consistent with the notion that the ancestral function of PCSK1 and PCSK2 in Bilateria is in neurosecretion. Further, our results are consistent with other data that, although PCSK1 and PCSK2 are generally expressed in the neural tissue, they can be occasionally recruited to other tissue types, such as exocrine glands, in some evolutionary lineages.

Our molecular phylogeny data suggested that PCSK1 and PCSK2 are not sister groups (Figure 4D), and therefore, this phylogenetic pattern would suggest independent recruitment of PCSK1 and PCSK2 to the neural and neuroendocrine functions or loss of these functions from other OGs. Unfortunately, the currently available data are not sufficient to resolve the detailed early evolutionary history of PCSK1 and PCSK2 OGs, or of the cell types that specifically express these genes. Further expansion of taxon sampling may help to explore the origins of the peptidergic nervous system and the peptide hormone system.

#### Widespread expression of PCSK7, PCSKX, Furin1, and Furin2

Consistent with our prediction, we found that furin-like PCSKs, namely *Hau-furin1.1*, *Hau-furin1.2*, *Hau-furin1.3*, *Hau-furin1.4*, *Hau-furin1.5*, *Hau-furin1.6*, and *Hau-furin2-like*, were broadly expressed (Figure 5F-5K and 5M), although our data suggest the possibility that some these genes are differentially expressed as well (*c.f.*, Figure 5G and 5H). Likewise, *Hau-pcsk7* and *Hau-pcskx* were also broadly expressed (Figure 5N, 5O). In contrast, the expression of *Hau-furin1.7* is restricted to the rear sucker (Figure 5L). These data suggested that genes encoding leech PCSK7, PCSKX, and all but one furin-like convertase exhibit far less tissue-type specificity in their expression than do PCSK1 and PCSK2. Our data also revealed that PCSKX is similar to PCSK7 and furin-like in exhibiting broad expression. We speculate that functional differences among these broadly expressed proprotein convertases may lie in their substrate preferences and/or in their subcellular localization, but testing this idea is beyond the scope of this study. If the absence of adult tissue-type specialization for these genes is conserved among bilaterians, it would suggest that tissue-specific expression patterns of mammalian PCSK4 and *Hau-pcsk1.7* may reflect neo-functionalization after gene duplication.

### Expression patterns of proprotein convertase genes in DV patterning of the germinal band

A key function of furin and PCSK5-7 in vertebrates is to process the developmentally important BMP ligands by removing the prodomain (Dubois, et al. 1995; Cui, et al. 1998; Constam and Robertson 2000; Essalmani, et al. 2008; Nelsen and Christian 2009). Although they are broadly expressed in adult tissue, selected proprotein convertase genes are upregulated in cells of the embryonic signaling centers that produce active BMP and other TGFβ ligands in vertebrates (Constam, et al. 1996; Nelsen, et al. 2005). To see if this phenomenon is conserved between the leech and vertebrates, we next examined the expression patterns of the eight furin-like PCSKs, PCSK7 and PCSKX in stage-8 embryos of *Helobdella*. In this stage, BMP genes are expressed and are involved in dorsoventral patterning of the trunk ectoderm (Kuo and Weisblat 2011b), and we focused on testing if any of these proprotein convertases is upregulated in the critical BMP-expressing cells in the dorsoventral patterning process.

In the leech, ectoderm and mesoderm of the segmented trunk arise from five bilateral pairs of teloblast stem cells (Weisblat and Shankland 1985). These five pairs of teloblasts arise from the D quadrant of the four-cell embryo through a stereotypic series of asymmetric cell divisions (Figure 6A). The M teloblasts give rise to the mesoderm, and the N, O/P, O/P and Q teloblasts to the ectoderm. Each teloblast undergoes a repetitive series of asymmetric cell division to produce a bandlet of blast cells, each of which develops in a lineage-specific manner and gives rise to a definitive set of differentiated cells. In stage 8, the five bandlets from the five ipsilateral teloblasts converge at the posterior end of the embryo to form left and right germinal bands, respectively (Figure 6B). Within each germinal band, the four ectodermal bandlets form a parallel array superficial to the mesodermal bandlet. The dorsalmost ectodermal bandlet, the q bandlet, expresses BMP5-8 (Figure 6C), which induces the dorsolateral, P fate in cells of the adjacent bandlet produced by one of the O/P teloblasts (Kuo and Weisblat 2011b). Cells adopt the ventrolateral, O fate in the other O/P-derived bandlet that is not in direct contact with the q bandlet (Huang and Weisblat 1996; Kuo and Weisblat 2011b). Fates of the ventralmost ectodermal bandlet, the n bandlet, are not regulated by BMP5-8 signaling (Kuo and Weisblat 2011b). In addition to the q bandlet-specific BMP5-8 expression (Figure 6C), three other BMP genes (BMP2/4a, BMP2/4b, and ADMP) are broadly expressed in the germinal band (Kuo and Weisblat 2011b). The BMP2/4s also contribute to generating a dorsoventral gradient of BMP activity in the germinal band, but the function of ADMP is largely unknown (Kuo and Weisblat 2011b).

**Figure 6.**
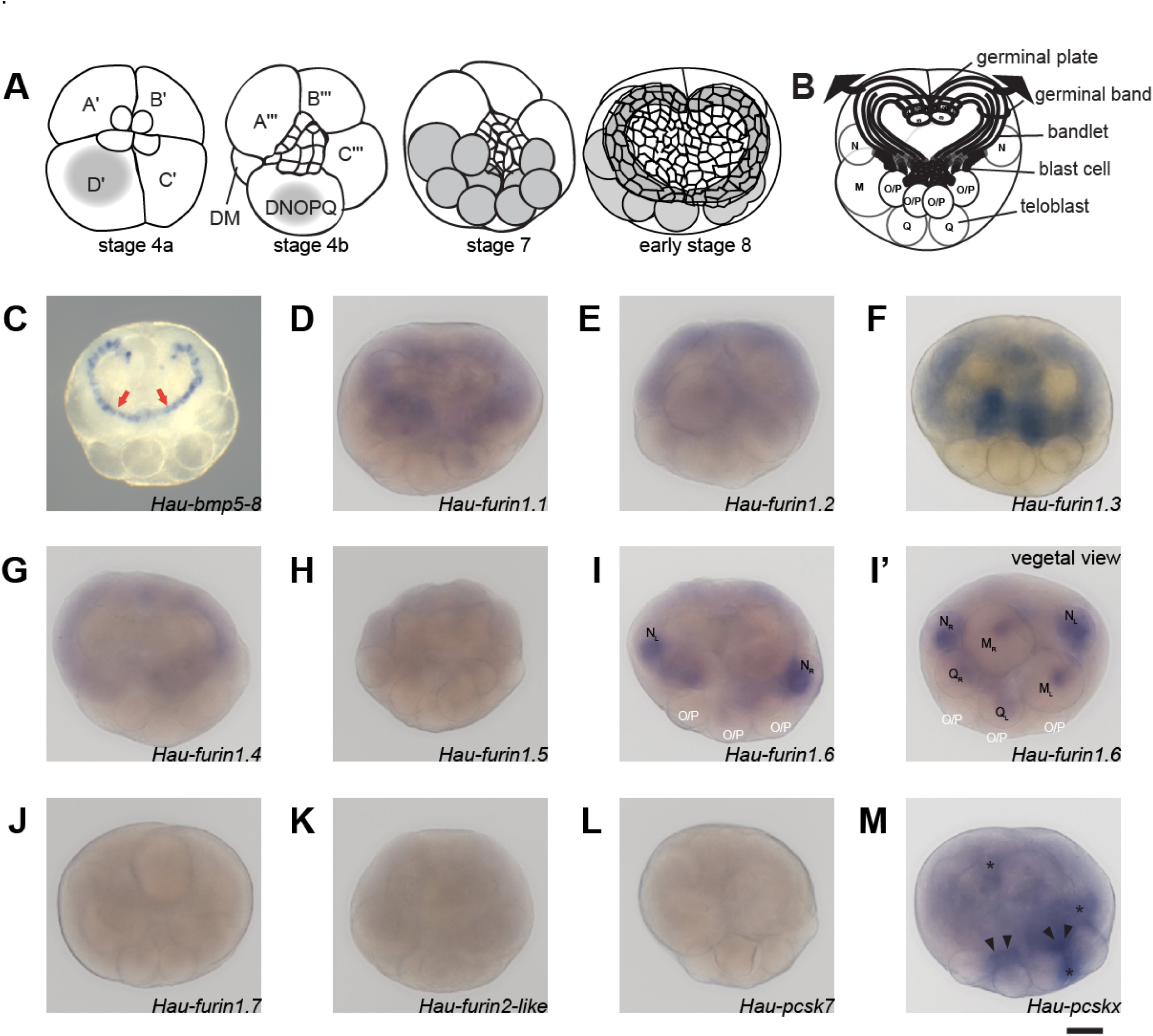
Expression of the Kexin-like genes in stage 8 *Helobdella austinensis* embryos revealed by whole-mount *in situ* hybridization. **A.** Developmental origin of teloblasts. Grey shade marks teloplasm, a special pool of cytoplasm that contains the teloblast determinants and is segregated into teloblasts during cleavage. **B.** Anatomical organization of the stage-8 leech embryo (see text for details). **C.** *Hau-bmp5-8* is specifically expressed in the dorsal, q bandlet. The image was taken from Kuo and Weisblat, 2011. **D-M.** Expression patterns of Kexin-like genes in stage-8 leech embryos. See the text for a detailed description. The asterisks in panel M mark the cytoplasm associated with macromere nuclei; the black arrowheads mark the isolated o/p bandlets entering the germinal bands. All panels expect I’ are in the animal-pole view; panel I’ is the vegetal-pole view. Scale bar = 100 μm.

Given the importance of the q bandlet in the BMP-mediated dorsoventral patterning in the leech, we predicted that one or more proprotein convertases would be upregulated in this region. In contrast to this prediction, however, the proprotein convertases, if expressed in the germinal band, showed a broad but weak expression throughout the germinal bands, and none exhibited an elevated expression level in the q bandlet (Figure 6D-M). For *Hau-furin1.1*, *Hau-furin1.2*, *Hau-furin1.3,* and *Hau-furin1.4*, the staining appeared to be somewhat stronger in the mesodermal bandlets (Figure 6D-6G). Staining for *Hau-furin1.5* was barely above the background level and was also stronger in the mesodermal bandlets (Figure 6H). *Hau-furin1.6* was expressed in all but the four O/P teloblasts (Figure 6I, 6I’). Expression of *Hau-furin1.7*, *Hau-furin2-like*, and *Hau-pcsk7* was not detected in stage 8 embryos (Figure 5J-L); this latter result was unlikely to arise from any technical issue since these probes did detect transcript expression, at even lower probe concentrations (1/5 of that used on the stage-8 embryo) when applied to juvenile specimens. *Hau-pcskx* transcript was somewhat elevated in the cytoplasm of macromeres as well as cells of the isolated o/p bandlets before they converge to form the germinal band (Figure 6M). Given that a proprotein convertase generally cleaves multiple substrates, these PCSKs may act on other molecules than BMPs. In any case, our observation implies that proteolytic processing of the BMP ligands may not have the same role in generating their axial patterning activity in the leech as in the vertebrates.

### Proteolytic processing by proprotein convertase is not required for the dorsalizing activity of Hau-BMP5-8

The low levels of proprotein convertase transcripts in the q bandlet suggest that levels of convertase activity are low in the germinal band tissue. If so, it could be (1) that cleavage by PCSK is irrelevant to the P fate-inducing activity of Hau-BMP5-8, or (2) that convertase-mediated cleavage is a rate-limiting step in P fate induction. To distinguish between these two alternatives, we sought to determine whether convertase-dependent proteolytic processing of Hau-BMP5-8 is required for its P fate-inducing activity, as assayed by the induction of *Hau-six1/2a* expression in the O-P equivalence group (Figure 7C). In BMPs and other members of the TGFβ family, the linker sequence between the prodomain and the bioactive TGFβ cysteine-knot domain contains at least one, often more than one, convertase cleavage site(s) (Derynck and Miyazono 2008). In Hau-BMP5-8, three such -RXXR- sites were identified, and we mutated all three sites into -GXXG- to prevent recognition and cleavage by convertases (Figure 7A). To see if this convertase-resistant Hau-BMP5-8 can induce P fate, we injected *in vitro* transcribed mRNA encoding the mutant Hau-BMP5-8 into the N teloblast and found that the o bandlet, which neighbors the n bandlet, was invariably induced to express the P fate marker, *Hau-six1/2a* ectopically (Figure 7E). This result was identical to that obtained from embryos in which wild-type Hau-BMP5-8 was expressed in the n bandlet (Figure 7D), indicating that proteolytic processing by proprotein convertase at the linker sequence is not required for the P fate-inducing activity of Hau-BMP5-8.

**Figure 7.**
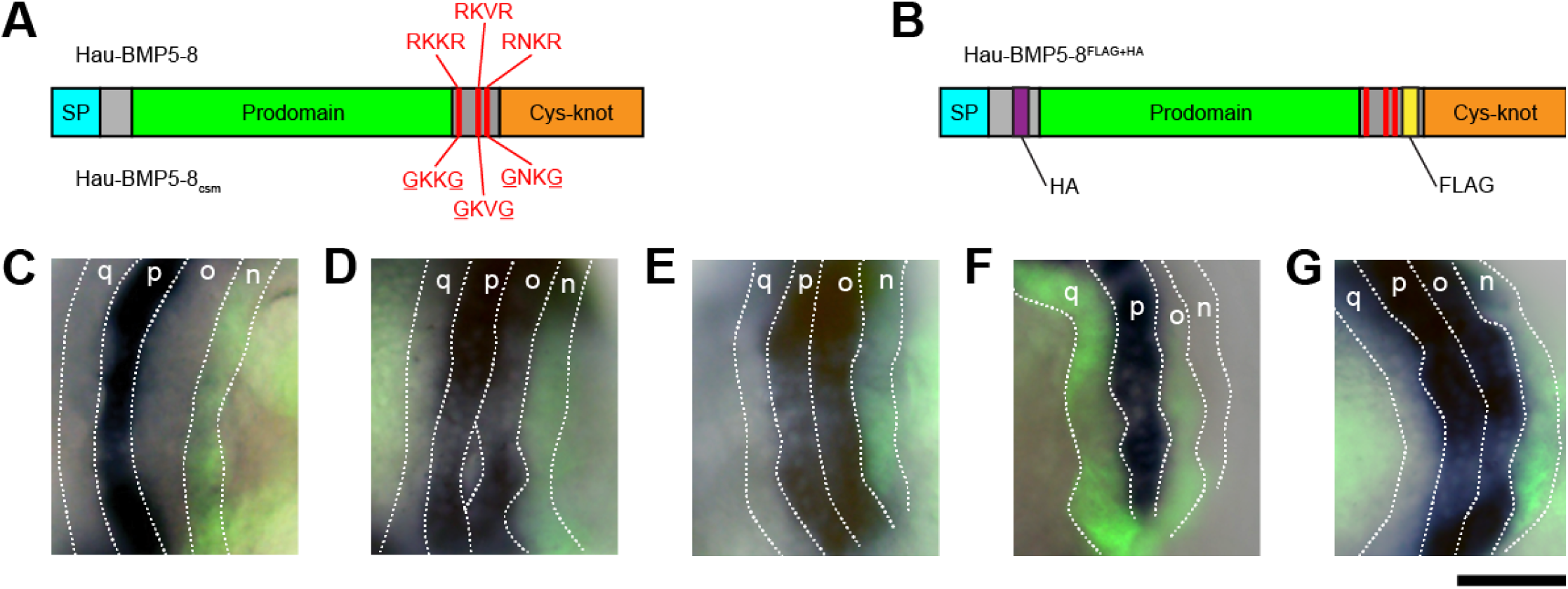
Cleavage by convertase is not required for the P fate-inducing activity of Hau-BMP5-8. **A.** Schematic representation of the wild-type Hau-BMP5-8 and Hau-BMP5-8_csm_, in which all three putative **c**onvertase **s**ites were **m**utated to make it resistant to convertase-mediated proteolytic cleavage. **B.** Schematic representation of HauBMP5-8^FLAG+HA^, in which a FLAG epitope tag was inserted between the TGFβ cysteine knot domain and the convertase cleavage site and an HA epitope tag was inserted between the signal peptide and the prodomain. In the mature, convertase-processed form, the HA tag should label the prodomain and the FLAG tag the active signaling ligand. **C.** Normal embryo in which the n bandlet was labeled with lineage tracer; the P fate marker *Hau-six1/2a* was only expressed in the p bandlet, as revealed by *in situ* hybridization. **D.** Embryo whose n bandlet mis-expressing the wild-type Hau-BMP5-8, *Hau-six1/2a* was ectopically expressed in the o bandlet, as well as the p bandlet. **E.** Similarly, *Hau-six1/2a* was also ectopically expressed in the o bandlet when the convertase-resistant Hau-BMP5-8_csm_ was mis-expressed in the n bandlet. **F.** When Hau-furin1.1 was mis-expressed in the o, p and q bandlets, the expression pattern of *Hau-six1/2a* remained normal, restricted to the p bandlet. No expansion of *Hau-six1/2a* expression to the o bandlet was observed. **G.** Hau-BMP5-8^FLAG+HA^ mis-expressed in the n bandlet induced ectopic *Hau-six1/2a* expression in the o bandlet as did wild-type Hau-BMP5-8 (D). Scale bar: 50 μm.

To further test the rate-limiting hypothesis, we next over-expressed the full-length wild-type Hau-furin1.1 in the germinal band by injecting its mRNA into a NOPQ proteloblast. In such embryos, no change in the expression of the P fate marker *Hau-six1/2a* was observed (Figure 7F). This result negated the hypothesis that convertase activity was rate-limiting for P-fate induction. Together, these results showed that proteolytic processing by convertase was neither necessary for P fate-inducing activity of Hau-BMP5-8, nor could it enhance BMP signaling in the leech germinal band.

On the other hand, the presence of proprotein convertase, even at a background level, should cleave at least a portion of the Hau-BMP5-8 protein. In an attempt to verify the cleavage of Hau-BMP5-8 in the germinal band, we generated a tagged Hau-BMP5-8 construct, in which an HA tag was inserted to the N-terminal region of the prodomain, and a FLAG tag was placed downstream to the convertase sites at the N-terminus of the TGFβ cysteine-knot domain. Cleavage of Hau-BMP5-8 should result in a separation of the HA-tagged prodomain fragment from the FLAG-tagged TGFβ cysteine-knot domain fragment. This tagged Hau-BMP5-8 is biologically active since it is capable of inducing *Hau-six1/2a* expression in the o bandlet when ectopically expressed in the n bandlet (Figure 7G). We then performed Western blot using lysates of embryos expressing the tagged Hau-BMP5-8 in the q bandlet or in the entire ectoderm. Despite multiple trials under various conditions and using different antibodies, such Western blot experiments were not successful due to commercial antibodies’ prevalent cross-reactivities to endogenous leech antigens. We also replaced both HA and FLAG tags in the Hau-BMP5-8 construct with GFP, but unfortunately, the same technical issue persisted.

Convertase-mediated cleavage separates the bioactive TGFβ cysteine-knot domain from the prodomain (Panganiban, et al. 1990; Dubois, et al. 1995; Cui, et al. 1998; Birsoy, et al. 2005), and it has been shown that association with prodomain may bring about different consequences for different TGFβ-family ligands, in different species, and in different cellular environments: in some cases, the prodomain anchors the TGFβ-family ligand to the extracellular matrix; in others, the prodomain prevents the ligand from binding its receptor; in yet other cases, the prodomain has no effect on signaling activity of the ligand (Gregory, et al. 2005; Sengle, et al. 2011; Jiang, et al. 2016). Although we had not succeeded in verifying the cleavage of Hau-BMP5-8 in the germinal band, we nevertheless demonstrated that such cleavage is irrelevant to Hau-BMP5-8’s ability to induce the P fate. This finding is in stark contrast to the situation in vertebrate embryos, in which overexpression of BMP4 or BMP7 with mutated cleavage sites has a dominant-negative effect (Hawley, et al. 1995); on the other hand the apparent irrelevance of convertase-mediated cleavage in the leech germinal band resembles the convertase-resistant mutant of the zebrafish nodal homolog Southpaw, which exhibits a similarly short-range signal activity, rather than a complete loss of signal (Tessadori, et al. 2015). The endogenous Hau-BMP5-8 may likewise be in a prodomain-bound form, regardless of whether it is proteolytically processed or not, and the presence of the prodomain may restrict the signaling range of BMP5-8, without interfering with its receptor binding.

Although technical difficulties prevented us from demonstrating the cleavage of Hau-BMP5-8 directly, finding that over-expression of proprotein convertases did not expand the signal range of Hau-BMP5-8 suggests that its prodomain and TGFβ cysteine-knot domain may have remained in association even after being cleaved. This would explain the convertase-independent, short-range signaling activity in the q bandlet and for the absence of a lineage-specific expression of proprotein convertase. We note that the *Drosophila* Gbb ligand, also a BMP5-8 homolog, has a stronger signaling activity and a longer signaling range when covalently linked to the C-terminal portion of its prodomain than as the fully processed short ligand (Akiyama, et al. 2012). In contrast to Gbb, however, the unprocessed Hau-BMP5-8 would contain the complete prodomain because it lacks the homologous cleavage site within its prodomain; this may account for the different behaviors of these two BMP5-8 ligands. In any case, together, these cases have revealed diversity in the regulation of how the BMP ligands are distributed in extracellular space and the differing role of proprotein convertases in regulating the signaling range of BMP ligands in different biological contexts.

### Conclusions

Proprotein convertases are a well-studied group of endoproteases that have multiple roles in animal development and physiology. Here, we presented a large-scale phylogenetic survey of proprotein convertases across metazoan species. Our general findings were: (1) that there are six ancestral bilaterian OGs of Kexin-like convertases, including the previously unknown PCSKX; (2) spiralians, in general, have retained all six OGs while specific OGs were apparently lost in other lineages.

We have also used the leech *Helobdella* as a tractable spiralian model in which to carry out to begin a functional characterization of convertase OGs. Our results show that functional divergence among the PCSK OGs was probably already in place in the bilaterian ancestor. In contrast to the absolute requirement for convertase processing of BMP ligands in vertebrate embryos, we demonstrate an instance of short-range, convertase-independent BMP signaling in dorsoventral patterning of the leech embryo. Overall, our findings highlight the somewhat overlooked significance of such conserved “housekeeping” enzymes in the diversification of complex biological signaling systems and illustrate the importance of studying non-model organisms for illuminating the breadth of developmental mechanisms.

## Methods

### Gene Identification and Phylogenetic Analysis

The *Helobdella* proprotein convertase genes were identified by querying the filtered model protein dataset in the JGI *Helobdella robusta* database (https://genome.jgi.doe.gov/Helro1/Helro1.home.html) for proteins containing the Peptidase_S8 domain (Pfam: PF00082) and by a BLASTP search against using human furin as a query. The results from these two searches were compared, and the retrieved sequences were analyzed using HMMER to evaluate the completeness of the gene models. The incomplete models were manually curated by referencing the sequences of the isolated cDNA clones.

For phylogenetic reconstruction, gene models from other species were collected by querying the Ensembl genome databases. The amphioxus genome is not included in the Ensembl collection, and we queried the JGI *Branchiostoma floridae* database (https://genome.jgi.doe.gov/Brafl1/Brafl1.home.html) instead. Models missing any of the defining criteria were further analyzed by BLASTP search against the GenBank to identify the false positives and the incomplete models. Incomplete models were included in the gene copy analysis but were excluded from the phylogenetic analysis. Descriptive statistical data were processed and plotted using R packages. Conceptually translated amino acid sequences were aligned using MUSCLE (Edgar 2004b, a). RAxML (Stamatakis 2014) was used to produce Maximum Likelihood trees. Phylogenetic trees and statistical plots were edited to improve visual presentation by using Adobe Illustrator CS6.

### Molecular Cloning of *Helobdella* proprotein convertases genes

DNA fragments corresponding to the coding region of the *Helobdella* proprotein convertase genes were PCR amplified from a pool of mix-stage *H. austinensis* cDNA. *H. austinensis* is a sibling species of *H. robusta* and is more ready to breed and easier to maintain in the laboratory environment (Kutschera, et al. 2013). All experiments described here were performed with *H. austinensis* instead of *H. robusta*. PCR primers (Table S2) were designed using sequence information uncovered from the *H. robusta* genome database, and PCR reaction was performed using the proof-reading thermostable DNA polymerase Phusion (ThermoFisher). The obtained cDNA fragments were inserted into pGEM-T vector (Promega) by following standard cloning procedures. The cDNA clones were isolated and sequenced, and the sequence information was used to correct and supplement the gene models in the JGI *H. robusta* database, in case of mistaken annotation.

### Whole-Mount *In Situ* Hybridization

Linear DNA templates for probe synthesis were PCR amplified from the cDNA clones in the pGEM-T vector using SP6 and T7 primers and purified by gel extraction. Digoxigenin labeled antisense probes were *in vitro* transcribed with either SP6 or T7 RNA polymerase (ThermoFisher), depending on the orientation of cDNA insertion in the vector. Probe synthesis and whole-mount *in situ* hybridization were performed as previously described (Weisblat and Kuo 2009). The color reaction was carried out by using BM Purple alkaline phosphatase substrate (Roche). Following the color reaction, the specimens were stored in ethanol and were mounted in 70% glycerol solution for imaging. Digital images were obtained by using an Olympus DP80 CCD camera mounted on an Olympus BX63 microscope. The acquired images were adjusted for size, contrast, and brightness by using Adobe Photoshop CS6, and the adjusted images were assembled into figure panels and annotated using Adobe Illustrator CS6.

### Expression of Molecular Constructs

Two different approaches were used to express Hau-BMP5-8 variants in the specified cell lineage. In one, synthesized capped RNA was injected into a teloblast or proteloblast (Zhang and Weisblat 2005); alternatively, plasmid DNA carrying the leech EF1α upstream sequence that is capable of driving transgene expression (Pilon and Weisblat 1997) was injected instead. These two methods yielded similar results. All experiments were performed with a >30 sample size; similar to the previously published Hau-BMP5-8 mis-expressing experiments (Kuo and Weisblat 2011b), invariable results were obtained in each set of experiments

Capped RNA was *in vitro* transcribed from linearized plasmid template using mMESSAGE mMACHINE SP6 Kit (ThermoFisher) and was further polyadenylated by using Poly(A) Tailing Kit (ThermoFisher). The plasmid template for the wild-type Hau-BMP5-8, pCS107-Hau-bmp5-8, and the dosage of mRNA injection was also the same as previously described (Kuo and Weisblat 2011b). The Hau-BMP5-8 variants with epitope tag insertion and/or site-directed mutagenesis were generated by inverse PCR from the wild-type pCS107-Hau-bmp5-8 plasmid template, using a previously described strategy (Fisher and Pei 1997). The proofreading thermophilic DNA polymerase Phusion (ThermoFisher) was used in these PCR reactions. Constructs with multiple tags and/or mutations were produced by stepwise introduction of individual insertions and mutations. The primer pairs that were used to generate a specific construct were listed in Table S3.

The coding region of each Hau-BMP5-8 variant was then subcloned from the pCS107 plasmid into a site immediately downstream to the leech EF1α upstream sequence in pMiEF1P, which was modified from a Minos transposon vector (Pavlopoulos and Averof 2005). Plasmid DNA was injected at a concentration of 100 ng/μl in micropipette as previously described (Gline, et al. 2009).

Overexpression of Hau-furin1.1 was performed by plasmid injection. The full-length coding region cDNA of Hau-furin1.1 was PCR amplified by using a primer pair listed in Table S3 and then inserted into the pMiEF1P to produce the expression plasmid pMiEF1P-Hau-furin1.1.

## Supporting information

Supplemental Tabels

## Acknowledgments

This work was supported by a grant from MOST, Taiwan (102-2311-B-002-024-MY2) to DHK and NIH grant 2R01GM074619 to DAW.

