## Supplemental Tabels for "Diversification of metazoan Kexin-like proprotein convertases: insights from the leech *Helobdella*"

Table S1 List of gene models surveyed in this study

| Species (Database) | Gene ID | Gene Name | Label in Figs.2&3 | Gene Family |
|---|---|---|---|---|
| *Saccharomyces cerevisiae* (ENSEMBL) | YNL238W | *KEX2* | Sce-kexin | Kexin-like |
| | YCR045C | *RRT12* | | PK-like |
| | YEL060C | *PRB1* | | PK-like |
| | YOR003W | *YSP3* | | PK-like |
| *Acanthamoeba castellanii* (ENSEMBL) | ACA1_057280 | | | Kexin-like |
| | ACA1_234230 | | | Kexin-like |
| | ACA1_387910 | | | Kexin-like |
| | ACA1_264710 | | | S1P-like |
| | ACA1_222700 | | | PK-like |
| | ACA1_128830 | | | PK-like |
| | ACA1_223330 | | | PK-like |
| | ACA1_273880 | | | PK-like |
| | ACA1_321400 | | | PK-like |
| | ACA1_155550 | | | PK-like |
| | ACA1_278460 | | | PK-like |
| | ACA1_316750 | | | PK-like |
| | ACA1_326080 | | | PK-like |
| | ACA1_149240 | | | TPPII |
| *Capsaspora owczarzaki* (ENSEMBL) | CAOG_004148 | | Cow-pcsk | Kexin-like |
| | CAOG_005995 | | | S1P-like |
| | CAOG_003679 | | | PK-like |
| | CAOG_006156 | | | PK-like |
| | CAOG_007483 | | | TPPII |
| *Monosiga brevicollis* (ENSEMBL) | MONBRDRAFT_30798 | | Mbr-30798 | Kexin-like |
| | MONBRDRAFT_14515 | | Mbr-14515 | Kexin-like |
| | MONBRDRAFT_27758 | | | S1P-like |
| | MONBRDRAFT_34612 | | | TPPII |
| *Amphimedon queenslandica* (ENSEMBL) | Aqu2.1.25568 | | Aqu-25568 | Kexin-like |
| | Aqu2.1.31170 | | Aqu-31170 | Kexin-like |
| | Aqu2.1.28225 | | Aqu-28225 | Kexin-like |
| | Aqu2.1.24525 | | Aqu-24525 | Kexin-like |
| | Aqu2.1.31171 | | Aqu-31171 | Kexin-like |
| | Aqu2.1.23420 | | Aqu-23420 | Kexin-like |
| | Aqu2.1.11050 + Aqu2.1.11051 | | | Kexin-like |
| | Aqu2.1.05739 | | | Kexin-like |
| | Aqu2.1.11049 | | | Kexin-like |
| | Aqu2.1.44231 | | | S1P-like |
| | Aqu2.1.38120 | | | PK-like |
| | Aqu2.1.39723 | | | PK-like |
| | Aqu2.1.38118 | | | PK-like |
| | Aqu2.1.37599 | | | PK-like |
| | Aqu2.1.38117 | | | PK-like |
| | Aqu2.1.32598 | | | PK-like |

| Species (Database) | Gene ID | Gene Name | Label in Figs.2&3 | Gene Family |
|---|---|---|---|---|
| *Amphimedon queenslandica* (ENSEMBL) cont. | Aqu2.1.30105 | | | PK-like |
| | Aqu2.1.38114 | | | PK-like |
| | Aqu2.1.38119 | | | PK-like |
| | Aqu2.1.38116 | | | PK-like |
| | Aqu2.1.08606 + Aqu2.1.08607 | | | PK-like |
| | Aqu2.1.00245 | | | PK-like |
| | Aqu2.1.30109 | | | PK-like |
| | Aqu2.1.38115 | | | PK-like |
| | Aqu2.1.02987 | | | TPPII |
| | Aqu2.1.43819 | | | TPPII |
| *Mnemiopsis leidyi* (ENSEMBL) | ML11682a | | Mle-11682 | Kexin-like |
| | ML20261a | | Mle-20261 | Kexin-like |
| | ML207912a | | Mle-207912 | Kexin-like |
| | ML07022a | | Mle-7022 | Kexin-like |
| | ML200226a | | Mle-200226 | Kexin-like |
| | ML279823a | | Mle-279823 | Kexin-like |
| | ML009145a | | | S1P-like |
| *Trichoplax adherens* (ENSEMBL) | TriadG60295 | | Tad-60295 | Kexin-like |
| | TriadG22925 | | Tad-22925 | Kexin-like |
| | TriadG54612 | | Tad-54612 | Kexin-like |
| | TriadG19704 | | | S1P-like |
| | TriadG61672 | | | TPPII |
| *Nematostella vectensis* (ENSEMBL) | NEMVEDRAFT_v1g234208 | | Nve-234208 | Kexin-like |
| | NEMVEDRAFT_v1g348 | | Nve-348 | Kexin-like |
| | NEMVEDRAFT_v1g129977 | | Nve-129977 | Kexin-like |
| | NEMVEDRAFT_v1g161314 | | Nve-161314 | Kexin-like |
| | NEMVEDRAFT_v1g160244 | | Nve-160244 | Kexin-like |
| | NEMVEDRAFT_v1g112522 | | Nve-112522 | Kexin-like |
| | NEMVEDRAFT_v1g168742 | | Nve-168742 | Kexin-like |
| | NEMVEDRAFT_v1g134107 + NEMVEDRAFT_v1g2207 | | Nve-184 | Kexin-like |
| | NEMVEDRAFT_v1g112587 + NEMVEDRAFT_v1g112538 | | | Kexin-like |
| | NEMVEDRAFT_v1g225607 | | | S1P-like |
| | NEMVEDRAFT_v1g225649 | | | S1P-like |
| | NEMVEDRAFT_v1g225649 | | | TPPII |
| *Lottia gigantean* (ENSEMBL) | LotgiG106119 | | Lgi-106119 | Kexin-like |
| | LotgiG81964 | | Lgi-81964 | Kexin-like |
| | LotgiG102960 | | Lgi-102960 | Kexin-like |
| | LotgiG212426 | | Lgi-212426 | Kexin-like |
| | LotgiG112524 | | Lgi-112524 | Kexin-like |
| | LotgiG224835 | | Lgi-224835 | Kexin-like |
| | LotgiG123675 | | | S1P-like |
| | LotgiG129137 | | | TPPII |
| *Crassostrea gigas* (ENSEMBL) | CGI_10016370 | | Cgi-34782 | Kexin-like |

| Species (Database) | Gene ID | Gene Name | Label in Figs.2&3 | Gene Family |
|---|---|---|---|---|
| *Crassostrea gigas* (ENSEMBL) cont. | CGI_10003367 | | Cgi-26985 | Kexin-like |
| | CGI_10003191 | | Cgi-21858 | Kexin-like |
| | CGI_10019653 | | Cgi-23284 | Kexin-like |
| | CGI_10022584 | | Cgi-35453 | Kexin-like |
| | CGI_10027463 | | Cgi-29248 | Kexin-like |
| | CGI_10013398 | | Cgi-25267 | Kexin-like |
| | CGI_10004432 | | | Kexin-like |
| | CGI_10009470 | | | S1P-like |
| | CGI_10005825 | | | TPPII |
| *Octopus bimaculoides* (ENSEMBL) | Ocbimv22002644m.g | | Obi-02644 | Kexin-like |
| | Ocbimv22020256m.g | | Obi-20256 | Kexin-like |
| | Ocbimv22016958m.g | | Obi-16958 | Kexin-like |
| | Ocbimv22028944m.g | | Obi-28944 | Kexin-like |
| | Ocbimv22033366m.g | | | Kexin-like |
| | Ocbimv22014564m.g | | | S1P-like |
| | Ocbimv22020590m.g | | | TPPII |
| *Capitella teleta* (ENSEMBL) | CapteG52595 | | Cte-52595 | Kexin-like |
| | CapteG161485 | | Cte-161485 | Kexin-like |
| | CapteG153069 | | Cte-153069 | Kexin-like |
| | CapteG121853 | | Cte-121853 | Kexin-like |
| | CapteG194857 | | Cte-194857 | Kexin-like |
| | CapteG177803 | | Cte-177803 | Kexin-like |
| | CapteG228241 | | | S1P-like |
| | CapteG217313 | | | PK-like |
| | CapteG169944 | | | PK-like |
| | CapteG220467 | | | PK-like |
| | CapteG54371 | | | PK-like |
| | CapteG127412 | | | PK-like |
| | CapteG54339 | | | PK-like |
| | CapteG83747 | | | PK-like |
| | CapteG121340 | | | PK-like |
| | CapteG61455 | | | PK-like |
| | CapteG184079 | | | PK-like |
| | CapteG163820 | | | PK-like |
| | CapteG217316 | | | PK-like |
| | CapteG228048 | | | TPPII |
| *Helobdella robusta* (ENSEMBL) | HelroG193616 | *Hro-pcsk1* | Hro-PCSK1 | Kexin-like |
| | HelroG96355 | *Hro-pcsk2* | Hro-PCSK2 | Kexin-like |
| | HelroG84740 | *Hro-furin1.1* | Hro-furin1.1 | Kexin-like |
| | HelroG170307 | *Hro-furin1.2* | Hro-furin1.2 | Kexin-like |
| | HelroG107946 | *Hro-furin1.3* | Hro-furin1.3 | Kexin-like |
| | HelroG82350 | *Hro-furin1.4* | Hro-furin1.4 | Kexin-like |
| | HelroG105284 | *Hro-furin1.5* | Hro-furin1.5 | Kexin-like |
| | HelroG190143 + HelroG93024 | *Hro-furin1.6* | Hro-furin1.6 | Kexin-like |
| | HelroG72480 | *Hro-furin1.7* | Hro-furin1.7 | Kexin-like |
| | HelroG74170 + Helro149650 + Helro137839 | *Hro-furin2-like* | Hro-furin2-like | Kexin-like |

| Species (Database) | Gene ID | Gene Name | Label in Figs.2&3 | Gene Family |
|---|---|---|---|---|
| *Helobdella robusta* (ENSEMBL) cont. | HelroG81814 + Helro184031 | *Hro-pcsk7* | Hro-PCSK7 | Kexin-like |
| | HelroG82977 | *Hro-pcskx* | Hro-PCSKX | Kexin-like |
| | HelroG194246 | | | S1P-like |
| | HelroG107485 | | | TPPII |
| *Lingula anatine* (ENSEMBL) | g1247 | | Lan-1247 | Kexin-like |
| | g33407 | | Lan-33407 | Kexin-like |
| | g4306 | | Lan-4306 | Kexin-like |
| | g15382 | | Lan-15382 | Kexin-like |
| | g404 | | Lan-404 | Kexin-like |
| | g15960 | | Lan-15960 | Kexin-like |
| | g6253 | | Lan-6253 | Kexin-like |
| | g405 | | Lan-405 | Kexin-like |
| | g1500 | | Lan-1500 | Kexin-like |
| | g27514 | | Lan-27514 | Kexin-like |
| | g8872 | | Lan-8872 | Kexin-like |
| | g25567 | | Lan-25567 | Kexin-like |
| | g4660 + g4661 | | | Kexin-like |
| | g24092 + g24091 | | | Kexin-like |
| | g2559 + g2560 | | | Kexin-like |
| | g26156 | | | Kexin-like |
| | g11440 | | | Kexin-like |
| | g31800 | | | S1P-like |
| | g8392 | | | S1P-like |
| | g6563 + g6562 | | | S1P-like |
| | g20349 | | | PK-like |
| | g19508 | | | PK-like |
| | g20348 | | | PK-like |
| | g33287 | | | PK-like |
| | g10243 | | | PK-like |
| | g11146 | | | PK-like |
| | g11147 | | | PK-like |
| | g14553 | | | PK-like |
| | g10176 | | | PK-like |
| | g14555 | | | PK-like |
| | g14554 | | | PK-like |
| | g19487 | | | PK-like |
| | g17482 | | | PK-like |
| | g12444 | | | PK-like |
| | g22139 | | | PK-like |
| | g26421 | | | PK-like |
| | g33111 | | | PK-like |
| | g10242 | | | PK-like |
| | g10384 | | | PK-like |
| | g13429 | | | PK-like |
| | g32391 | | | PK-like |
| | g10239 | | | PK-like |
| | g10385 | | | PK-like |
| | g12204 | | | PK-like |

| Species (Database) | Gene ID | Gene Name | Label in Figs.2&3 | Gene Family |
|---|---|---|---|---|
| *Lingula anatine* (ENSEMBL) cont. | g22137 | | | PK-like |
| | g16160 | | | PK-like |
| | g19488 | | | PK-like |
| | g32390 | | | PK-like |
| | g10241 | | | PK-like |
| | g31346 | | | PK-like |
| | g13431 | | | PK-like |
| | g12443 | | | PK-like |
| | g12202 | | | PK-like |
| | g12201 | | | PK-like |
| | g16771 | | | PK-like |
| | g19485 | | | PK-like |
| | g5629 | | | PK-like |
| | g3976 | | | PK-like |
| | g755 | | | PK-like |
| | g31345 | | | PK-like |
| | g3977 | | | PK-like |
| | g22552 | | | PK-like |
| | g4081 | | | PK-like |
| | g22549 | | | PK-like |
| | g10565 | | | PK-like |
| | g15680 | | | PK-like |
| | g5630 | | | PK-like |
| | g31844 | | | PK-like |
| | g26421 | | | PK-like |
| | g29291 | | | PK-like |
| | g31805 | | | PK-like |
| | g19878 | | | TPPII |
| | g30441 | | | TPPII |
| *Adineta vaga* (ENSEMBL) | GSADVT00036980001 | | Ava-36980 | Kexin-like |
| | GSADVT00007527001 | | Ava-07527 | Kexin-like |
| | GSADVT00040037001 | | Ava-40037 | Kexin-like |
| | GSADVT00045121001 | | Ava-45121 | Kexin-like |
| | GSADVT00031184001 | | Ava-31184 | Kexin-like |
| | GSADVT00040267001 | | Ava-40267 | Kexin-like |
| | GSADVT00059080001 | | Ava-59080 | Kexin-like |
| | GSADVT00030251001 | | Ava-30251 | Kexin-like |
| | GSADVT00046357001 | | Ava-46357 | Kexin-like |
| | GSADVT00002479001 | | Ava-02479 | Kexin-like |
| | GSADVT00004015001 | | Ava-04015 | Kexin-like |
| | GSADVT00048266001 | | Ava-48266 | Kexin-like |
| | GSADVT00019939001 | | Ava-19939 | Kexin-like |
| | GSADVT00030411001 | | Ava-30411 | Kexin-like |
| | GSADVT00046798001 | | Ava-46798 | Kexin-like |
| | GSADVT00039465001 | | Ava-39465 | Kexin-like |
| | GSADVT00017404001 + GSADVT0001740300 | | | Kexin-like |
| | GSADVT00058030001 | | | Kexin-like |
| | GSADVT00049407001 | | | Kexin-like |

| Species (Database) | Gene ID | Gene Name | Label in Figs.2&3 | Gene Family |
|---|---|---|---|---|
| *Adineta vaga* (ENSEMBL) cont. | GSADVT00039914001 | | | S1P-like |
| | GSADVT00018118001 | | | S1P-like |
| | GSADVT00006009001 | | | TPPII |
| | GSADVT00061992001 | | | TPPII |
| *Drosophila melanogaster* (ENSEMBL) | amon | amon | Dme-amon | Kexin-like |
| | Fur1 | furin1 | Dme-furin1 | Kexin-like |
| | Fur2 | furin2 | Dme-furin2 | Kexin-like |
| | S1P | S1P | | S1P-like |
| | TppII | TppII | | TPPII |
| *Tribolium castaneum* (ENSEMBL) | TC004402 | | Tca-pcsk1 | Kexin-like |
| | TC009181 | | Tca-pcsk2 | Kexin-like |
| | TC010869 | | Tca-furin1 | Kexin-like |
| | TC031960 | | Tca-furin2 | Kexin-like |
| | TC002816 | | | S1P-like |
| | TC034512 | | | TPPII |
| *Daphnia pulex* (ENSEMBL) | DAPPUDRAFT_59868 | | Dpu-59868 | Kexin-like |
| | DAPPUDRAFT_126713 | | Dpu-126713 | Kexin-like |
| | DAPPUDRAFT_43136 | | Dpu-43136 | Kexin-like |
| | DAPPUDRAFT_193075 | | Dpu-193075 | Kexin-like |
| | DAPPUDRAFT_306271 | | Dpu-306271 | Kexin-like |
| | DAPPUDRAFT_306585 | | Dpu-306585 | Kexin-like |
| | DAPPUDRAFT_310948 | | | S1P-like |
| | DAPPUDRAFT_186919 | | | TPPII |
| *Caenorhabditis elegans* (ENSEMBL) | aex-5 | aex-5 | Cel-aex5 | Kexin-like |
| | egl-3 | egl-3 | Cel-pcsk2 | Kexin-like |
| | kpc-1 | kpc-1 | Cel-kpc1 | Kexin-like |
| | bli-4 | bli-4 | Cel-bli4 | Kexin-like |
| | tpp-2 | tpp-2 | | TPPII |
| *Strongylocentrotus purpuratus* (ENSEMBL) | SP-NEC2_1 | SP-NEC2_1 | Spu-027781 | Kexin-like |
| | SP-FURIN1 | SP-FURIN1 | Spu-010722 | Kexin-like |
| | SP-FURIN | SP-FURIN | Spu-002615 | Kexin-like |
| | SP-NEC2 | SP-NEC2 | Spu-023813 | Kexin-like |
| | SP-FURIN_1 | SP-FURIN_1 | Spu-028030 | Kexin-like |
| | SP-PCSK7 | SP-PCSK7 | | Kexin-like |
| | SP-FURIN_2 | SP-FURIN_2 | | Kexin-like |
| | SP-MBTPS1 | SP-MBTPS1 | | S1P-like |
| | SP-ASP_8 | | | PK-like |
| | SP-SLSP_3 / SP-ASPL_2 | | | PK-like |
| | SP-HYPP_305 | | | PK-like |
| | SP-ASP_12 | | | PK-like |
| | SP-ASP_16 / SPU_014264 / SP-SLSP_10 | | | PK-like |
| | SP-HYPP_1141 | | | PK-like |
| | SP-SLSP_26 | | | PK-like |
| | SP-ASP-4 | | | PK-like |

| Species (Database) | Gene ID | Gene Name | Label in Figs.2&3 | Gene Family |
|---|---|---|---|---|
| *Strongylocentrotus purpuratus* (ENSEMBL) cont. | SP-SLSP_27 | | | PK-like |
| | SP-ASP_7 / SP-SLSP_17 | | | PK-like |
| | SP-ASP | | | PK-like |
| | SP-ASP-3 | | | PK-like |
| | SP-ASP_11 | | | PK-like |
| | SP-SLSP_18 | | | PK-like |
| | SP-SLSP_12 | | | PK-like |
| | SP-SLSP_25 / SP-ASP-6 / SP-ASP-5 | | | PK-like |
| | SP-SERP | | | PK-like |
| | SP-SLSP_13 | | | PK-like |
| | SP-SLSP_20 | | | PK-like |
| | SP-SLSP_8 | | | PK-like |
| | SP-SERP-2 | | | PK-like |
| | SP-SLSP_4 | | | PK-like |
| | SP-ASP_14 | | | PK-like |
| | SP-SLSP_14 | | | PK-like |
| | SP-SLSP_15 | | | PK-like |
| | SP-SLSP_21 | | | PK-like |
| | SP-SLSP_22 | | | PK-like |
| | SP-SERP-3 | | | PK-like |
| | SP-SLSP_24 | | | PK-like |
| | SP-SLSP_6 | | | PK-like |
| | SPU_025143 / SPU_009043 | | | PK-like |
| | SP-SLSP_1 | | | PK-like |
| | SP-SLSP_11 | | | PK-like |
| | SP-ASP_13 / SP-ASP_15 | | | PK-like |
| | SP-ASP_10 | | | PK-like |
| | SP-SLSP_7 | | | PK-like |
| | SP-PPCL | | | PK-like |
| | SP-ASP_9 | | | PK-like |
| | SP-ASP-2 | | | PK-like |
| | SP-ASP_17 | | | PK-like |
| | SP-SLSP_5 | | | PK-like |
| | SP-SLSP | | | PK-like |
| | SP-SLSP_9 | | | PK-like |
| | SP-SLSP_16 | | | PK-like |
| | SP-ASP_1 | | | PK-like |
| | SP-SLSP_2 | | | PK-like |
| | SPU_003191 | | | PK-like |
| | SPU_018846 | | | PK-like |
| | SPU_025576 | | | PK-like |
| | SP-SLSP_19 | | | PK-like |
| | SP-PCSK9L | | | PK-like |
| | SP-ASPL_1 | | | PK-like |
| | SPU_015237 | | | PK-like |
| | SPU_003192 | | | PK-like |
| | SP-HYPP_3123 | | | PK-like |
| | SP-SLSP_23 | | | PK-like |
| | SPU_025326 | | | PK-like |

| Species (Database) | Gene ID | Gene Name | Label in Figs.2&3 | Gene Family |
|---|---|---|---|---|
| *Strongylocentrotus purpuratus* (ENSEMBL) cont. | SPU_027076 | | | PK-like |
| | SPU_022967 | | | PK-like |
| | SPU_014673 | | | PK-like |
| | SP-ALLGIF1 | | | PK-like |
| | SP-SLSPL / SPU_012623 | | | PK-like |
| | SP-TPP2 | | | TPPII |
| *Branchiostoma floridae* (JGI) | Brafl84610 | | Bfl-84610 | Kexin-like |
| | Brafl98086 | | Bfl-98086 | Kexin-like |
| | Brafl66271 | | Bfl-66271 | Kexin-like |
| | Brafl75530 | | Bfl-75530 | Kexin-like |
| | Brafl98844 | | Bfl-98844 | Kexin-like |
| | Brafl105744 | | Bfl-105744 | Kexin-like |
| | Brafl119857 | | Bfl-119857 | Kexin-like |
| | Brafl1121298 | | Bfl-1121298 | Kexin-like |
| | Brafl123943 | | Bfl-123943 | Kexin-like |
| | Brafl165115 | | Bfl-165115 | Kexin-like |
| | Brafl1165859 | | Bfl-1165859 | Kexin-like |
| | Brafl166302 | | Bfl-166302 | Kexin-like |
| | Brafl174236 | | Bfl-174236 | Kexin-like |
| | Brafl85770 | | Bfl-85770 | Kexin-like |
| | Brafl91496 | | Bfl-91496 | Kexin-like |
| | Brafl124316 | | Bfl-124316 | Kexin-like |
| | Brafl144492 | | Bfl-144492 | Kexin-like |
| | Brafl254769 | | Bfl-254769 | Kexin-like |
| | Brafl65099 | | | S1P-like |
| | Brafl81022 | | | PK-like |
| | Brafl102644 | | | PK-like |
| | Brafl129459 | | | PK-like |
| | Brafl118415 | | | PK-like |
| | Brafl130739 | | | PK-like |
| | Brafl107715 | | | PK-like |
| | Brafl265827 | | | PK-like |
| | Brafl131810 | | | PK-like |
| | Brafl130113 | | | PK-like |
| | Brafl121465 | | | PK-like |
| | Brafl121466 | | | PK-like |
| | Brafl119698 | | | TPPII |
| | Brafl119705 | | | TPPII |
| *Ciona intestinalis* (ENSEMBL) | ENSCING00000006411 | | Cin-219063 | Kexin-like |
| | ENSCINT00000012624 | | Cin-294743 | Kexin-like |
| | ENSCINT00000011797 | | Cin-292557 | Kexin-like |
| | ENSCINT00000010394 | | Cin-248784 | Kexin-like |
| | ENSCINT00000012580 | | Cin-270411 | Kexin-like |
| | ENSCINT00000035891 | | | S1P-like |
| | ENSCINT00000016248 | | | TPPII |
| *Petromyzon marinus* (ENSEMBL) | ENSPMAT00000009791 | | | Kexin-like |

| Species (Database) | Gene ID | Gene Name | Label in Figs.2&3 | Gene Family |
|---|---|---|---|---|
| *Petromyzon marinus* (ENSEMBL) cont. | ENSPMAT00000009884 | | | Kexin-like |
| | ENSPMAT00000009883 | | | Kexin-like |
| | ENSPMAT00000001746 | | | Kexin-like |
| | ENSPMAT00000001849 | | | S1P-like |
| *Danio rerio* (ENSEMBL) | pcsk1 | pcsk1 | | Kexin-like |
| | pcsk2 | pcsk2 | | Kexin-like |
| | furinA | furinA | | Kexin-like |
| | furinB | furinB | | Kexin-like |
| | pcsk5a | pcsk5a | | Kexin-like |
| | pcsk5b | pcsk5b | | Kexin-like |
| | pcsk6 | pcsk6 | | Kexin-like |
| | pcsk7 | pcsk7 | | Kexin-like |
| | mbtps1 | mbtps1 | | S1P-like |
| | pcsk9 | pcsk9 | | PK-like |
| | tpp2 | tpp2 | | TPPII |
| *Xenopus tropicalis* (ENSEMBL) | pcsk1 | pcsk1 | Xtr-pcsk1 | Kexin-like |
| | pcsk2 | pcsk2 | Xtr-pcsk2 | Kexin-like |
| | furin | furin | Xtr-furin | Kexin-like |
| | pcsk4 | pcsk4 | Xtr-pcsk4 | Kexin-like |
| | pcsk5 | pcsk5 | Xtr-pcsk5 | Kexin-like |
| | pcsk6 | pcsk6 | Xtr-pcsk6 | Kexin-like |
| | pcsk7 | pcsk7 | Xtr-pcsk7 | Kexin-like |
| | mbtps1 | mbtps1 | | S1P-like |
| | pcsk9 | pcsk9 | | PK-like |
| | tpp2 | tpp2 | | TPPII |
| *Gallus gallus* (ENSEMBL) | PCSK1 | PCSK1 | Gga-pcsk1 | Kexin-like |
| | PCSK2 | PCSK2 | Gga-pcsk2 | Kexin-like |
| | FURIN | FURIN | Gga-furin | Kexin-like |
| | PCSK4 | PCSK4 | Gga-pcsk4 | Kexin-like |
| | PCSK5 | PCSK5 | Gga-pcsk5 | Kexin-like |
| | PCSK6 | PCSK6 | Gga-pcsk6 | Kexin-like |
| | PCSK7 | PCSK7 | Gga-pcsk7 | Kexin-like |
| | MBTPS1 | MBTPS1 | | S1P-like |
| | TPP2 | TPP2 | | TPPII |
| *Homo sapiens* (ENSEMBL) | PCSK1 | PCSK1 | Hsa-pcsk1 | Kexin-like |
| | PCSK2 | PCSK2 | Hsa-pcsk2 | Kexin-like |
| | FURIN | FURIN | Hsa-furin | Kexin-like |
| | PCSK4 | PCSK4 | Hsa-pcsk4 | Kexin-like |
| | PCSK5 | PCSK5 | Hsa-pcsk5 | Kexin-like |
| | PCSK6 | PCSK6 | Hsa-pcsk6 | Kexin-like |
| | PCSK7 | PCSK7 | Hsa-pcsk7 | Kexin-like |
| | MBTPS1 | MBTPS1 | | S1P-like |
| | PCSK9 | PCSK9 | | PK-like |
| | TPP2 | TPP2 | | TPPII |

Table S2. PCR primers used to amplify cDNA fragments of the 12 *Helobdella austinensis* kexin-like convertase

| Primer IDs | Sequence (5’ to 3’) |
|---|---|
| Hau-pcsk1 forward | 5’-TGGACGAGGTGTCGTCGTCA-3’ |
| Hau-pcsk1 reverse | 5’-GGATGTTCTGTGTCTTTGCA-3’ |
| Hau-pcsk2 forward | 5’-CTTCGAACACAAACTCGTTCA-3’ |
| Hau-pcsk2 reverse | 5’-TCAAAACTCGCTAAACTTCCT-3’ |
| Hau-furin1.1 forward | 5’-CCGTCCAAATTGAGGGAGGAA-3’ |
| Hau-furin1.1 reverse | 5’-CATGAAAGCCCAGTTGGTGA-3’ |
| Hau-furin1.2 forward | 5’-CGCGAAATACATAGCGCGAAA-3’ |
| Hau-furin1.2 reverse | 5’-CAATTGGTGAAACCCTCGTT-3’ |
| Hau-furin1.3 forward | 5’-CAGATCGACGGCAATGAAACAGT-3’ |
| Hau-furin1.3 reverse | 5’-GCGCACATGCAAAACTATCT-3’ |
| Hau-furin1.4 forward | 5’-GCGAAAGAAGTTGCCAGGGA-3’ |
| Hau-furin1.4 reverse | 5’-CGTCGTCATGAAAGTCCAGT-3’ |
| Hau-furin1.5 forward | 5’-AGATCGCTTTCAAGCAACAGTTATCA-3’ |
| Hau-furin1.5 reverse | 5’-TCAAAATCGTCTTTGGCAAT-3’ |
| Hau-furin1.6 forward | 5’-CAGTCTGGTACAGGCATTCCTATTGGT-3’ |
| Hau-furin1.6 reverse | 5’-CAAGACATCATTCATGGTAATGA-3’ |
| Hau-furin1.7 forward | 5’-GTACAGAAATGAATGGGCAGT-3’ |
| Hau-furin1.7 reverse | 5’-CCATTCATTGAAACCTTCGTT-3’ |
| Hau-furin2-like forward | 5’-GGCAGTGGAAATTGAGGGTGGT-3’ |
| Hau-furin2-like reverse | 5’-CTTCCAGGTACCTCTGGGAT-3’ |
| Hau-pcsk7 forward | 5’-CATAACATCACTGGCAGGGGCA-3’ |
| Hau-pcsk7 reverse | 5’-GAAGGTATCGTAGAGTGCCAT-3’ |
| Hau-pcskx forward | 5’-CGGCCTGGGTCAAAGGTTACA-3’ |
| Hau-pcskx reverse | 5’-GATGGATTCTCACCCCAGCT-3’ |

Table S3. PCR primers for building constructs and performing site-direct mutagenesis

| Construct<br>Primer IDs | Sequence (5' to 3') |
|---|---|
| pCS107 Hau-bmp5-8$_{csm}$ | |
| Hau-bmp5-8 cleavage site 1 mutagenesis forward | 5'-GCCGACATAAGAAACCTCGGA-3' |
| Hau-bmp5-8 cleavage site 1 mutagenesis reverse | 5'-AGATCCTTTCTTTCCTCCACTCAACTGA-3' |
| Hau-bmp5-8 cleavage sites 2+3 mutagenesis forward | 5'-GATAGTGGGAACAAGGGAAATTCGTA-3' |
| Hau-bmp5-8 cleavage sites 2+3 mutagenesis reverse | 5'-CTTGCCCACTTTCCCGGTTTTGAA-3' |
| pCS107 Hau-bmp5-8$^{FLAG+HA}$ | |
| Hau-bmp5-8 FLAG insertion forward | 5'-GACGATAAGAAGAAACCAAACTATCTTACCAGACTCTGTCAAC-3' |
| Hau-bmp5-8 FLAG insertion reverse | 5'-GTCATCCTTGTAATCGTACGAATTTCTCTTGTTCCGACTATCCTTGC-3' |
| Hau-bmp5-8 HA insertion forward | 5'-GATGTTCCAGATTACGCTCAGCAGTATTCAAATGACGGCGGC-3' |
| Hau-bmp5-8 HA insertion reverse | 5'-GTATGGGTATTCGTGATGGTGATGTTGAGGTGGAG-3' |
| pCS107 Hau-bmp5-8$^{GFP}$ | |
| Hau-bmp5-8 NheI-SacI insertion forward | 5'-AGAGCTCTACAAGAAGAAACCAAACTATCTT-3' |
| Hau-bmp5-8 NheI-SacI insertion reverse | 5'-GGCTAGCCGAATTTCTCTTGTTCCGACTATC-3' |
| GFP forward + NheI site | 5'-GGCTAGCATGGTGAGCAAGGGCGAGGAG-3' |
| GFP reverse + SacI site | 5'-AGAGCTCCTTGTACAGCTCGTCCATGCCG-3' |
| pMiEF1P Hau-furin1.1 | |
| Hau-furin1.1 forward + ClaI site | GAACATCGATATGAATTTTATGATGATCATTTGCTACCT |
| Hau-furin1.1 reverse + XhoI site | GAACTCGAGAAATGCTAATAAATTTAAACAATAAAGATCATAATGA |
